# Differences in the structure of plant polygalacturonases specify enzymes’ dynamics and processivities to fine-tune cell wall pectins

**DOI:** 10.1101/2022.06.22.497136

**Authors:** Josip Safran, Wafae Tabi, Vanessa Ung, Adrien Lemaire, Olivier Habrylo, Julie Bouckaert, Maxime Rouffle, Aline Voxeur, Paula Pongrac, Solène Bassard, Roland Molinié, Jean-Xavier Fontaine, Serge Pilard, Corinne Pau-Roblot, Estelle Bonnin, Danaé Sonja Larsen, Mélanie Morel-Rouhier, Jean-Michel Girardet, Valérie Lefebvre, Fabien Sénéchal, Davide Mercadante, Jérôme Pelloux

## Abstract

The fine-tuning of pectins by polygalacturonases (PGs) plays a key role in modulating plant cell wall chemistry and mechanics, impacting plant development. In plants, the high number of PGs encoded in the genome questions the regulation of pectin depolymerization and the roles of distinct isozymes in the control of development. Here we report the first crystal structures of two PGs from Arabidopsis, PGLR and ADPG2 whose expression overlap in roots. Albeit having overall conserved folds and active sites, PGLR and ADPG2 differed in the structure of their binding grooves and in the amino-acids of the subsites. We determined the structural features that explain the absence of inhibition of the plant PGs by endogenous PG-Inhibiting Proteins (PGIPs). By combining molecular dynamic simulations, analysis of enzymes’ kinetics and hydrolysis products, we showed that subtle differences in PGLR and ADPG2 structures translated into distinct enzyme-substrate dynamics and enzymes’ processivities. Using the plant root as a developmental model, exogenous application of purified enzymes showed that these distinct PGLR/ADPG2 processivities ultimately translated into different impacts on development. The highly processive ADPG2 had major effects on both root cell elongation and cell adhesion. Our study suggests that, in plants, gene redundancy is unlikely to reflect redundant biochemical specificities. Isozymes of distinct specificities and processivities are likely to be of major importance for the fine spatial and temporal regulation of pectin structure.

**Significance Statement:** Plant polygalacturonases (PG) are enzymes that play a key role in the regulation of cell wall pectin chemistry by controlling the degree of polymerization of the HG chains. The high number of genes encoding PG in Arabidopsis questions the rationale for such abundance. We solved the crystal structure of two PG (PGLR and ADPG2) whose expression overlap in roots and showed, using combined computational and experimental approaches, that they differ in their enzyme-substrate dynamics, leading to distinct processivities. The highly processive ADPG2 can generate digestion products of shorter degree of polymerization, and upon exogenous application on developing roots, induced drastic developmental defects. Our study suggests that gene redundancy is unlikely to reflect redundant biochemical specificities of isozymes.

## Introduction

The plant primary cell wall, composed of an intricate network of polysaccharides and proteins, is constantly remodelled, translating in changes in its mechanical properties, which ultimately affect the extent of cell growth or the response to environmental stress (1). Pectin, the major polysaccharide of the primary cell wall of dicotyledonous species such as Arabidopsis, are composed of homogalacturonan (HG): a homopolymer of α-1,4-linked-D-galacturonic acid (GalA) units, that can be substituted with methylester and/or acetyl groups (2). The control of the degree of polymerization (DP) of HG by polygalacturonases (PGs) impacts diverse developmental processes such as root/hypocotyl growth, stomata functioning, cell separation during pollen formation and pollen tube elongation (3–8). Importantly, phytopathogenic organisms, including parasitic plants, also produce PGs, thus contributing to host colonization by degrading the physical barrier of the plant cell wall (9). Although all perform the hydrolysis of the α-(1–4) glycosidic bond between two adjacent non-methylesterified GalA units, PGs can differ in their mode of action and are referred to as endo-PGs (EC 3. 2.1.15) or exo-PGs (EC 3.2.1.67) if they either hydrolyse in the middle of the HG chain or attack from its non-reducing end (10, 11). All resolved structures of PGs from microorganisms fold into a right-handed parallel beta-helix and harbour four conserved amino acids (aa) stretches in their active site: namely NTD, DD, GHG and RIK (11). In a typical endo-PG, such as that from *Aspergillus aculeatus* PG1 (AaPG1), the active site is organized in a tunnel-like binding cleft, allowing the enzyme to bind the polysaccharide and produce pectic fragments called oligogalacturonides (OGs) of various DP and with different methyl and acetyl substitutions (12–14). In contrast, the structure of exo-PGs differs, loop extension turns the open-ended channel into a closed pocket, restricting the attack to the non-reducing end of the substrate, and releasing non-methylesterified GalA monomers or dimers (15). It has been reported that pathogenic PGs are inhibited by Polygalaturonase Inhibiting Proteins (PGIPs), expressed by plants upon infection, either through competitive or non-competitive interactions, in a strategic attempt by plants to limit pectin degradation and pathogenic invasion (16, 17). However, the nature of this inhibition is PG-specific as certain PGIPs were ineffective in mediating PG inhibition (17, 18). In contrast a number of plant PGs are not inhibited by plant PGIPs, which suggests yet unidentified and specific structural features among this class of enzymes. The PG-mediated degradation of HG can have two distinct consequences: i) it can impact polysaccharide rheology, decreasing cell wall stiffness and promoting cell growth (or infection by pathogens) and/or ii) it can produce OGs, which can act as signalling molecules (14, 19). It seems likely that the fine composition of OG arrays produced by a myriad of differentially expressed PG isoforms can modulate the oligosaccharide interactions with cell wall integrity receptors, triggering distinct downstream signalling events (13).

In plants, PGs are encoded by large multigenic families (68 genes in *Arabidopsis thaliana*), which questions the rationale for such an abundance in the context of the cell wall. Considering such a large number of genes, and potential compensation mechanisms mediated by partial functional redundancy between isoforms, the use of reverse or forward genetic mutants can only bring partial clues to sample the diversity of the plant PGs’ landscape.

Here we report on the biochemical and first structural characterization of two plant PGs, PGLR (PolyGalacturonase Lateral Root) and ADPG2 (Arabidopsis Dehiscence zone PolyGalacturonase 2), whose expression overlaps in Arabidopsis roots during lateral root emergence (8, 20–22). We first determined the structural features that explain the absence of inhibition of plant PG by endogenous PGIPs. We next found that, although having an overall conserved structure and overlapping functional profiles, enzymes have key and noticeable differences in their processivities. The investigation of PGLR and ADPG2 crystal structures, together with enzyme-substrate complexes, via combined experimental and computational approaches, including binding kinetics, Molecular Dynamics (MD) simulations, LC-MS/MS profiling of digestion products, indeed highlighted the existence of a link between enzyme-substrate interactions and dynamics, enzyme activities and processivities. Using exogenous application of purified enzymes as a tool, we further showed that these distinct modes of action can translate into peculiar effects on root development. This overall shows that, despite apparent gene redundancy, plant PGs have distinct biochemical activities leading to peculiar consequences on plant development, which could be a key for the fine, spatial and temporal, tuning of cell wall chemistry and mechanics.

## Results

### Crystal structures of *A. thaliana* PGLR and ADPG2 reveals conserved β-fold

ADPG2 was produced as an active recombinant proteins in the yeast *Pichia pastoris* and subsequently purified, similarly to PGLR (8). PGLR and ADPG2 present similar biochemical characteristics, with an optimal activity at acidic pH, temperatures ranging from 25 to 50 °C and on pectic substrates with low degree of methylesterification (DM, SI Appendix, Fig. S1 and (8)). Using PolyGalacturonic Acid (PGA) as a substrate, at 25°C, PGLR and ADPG2 differ in their Km (14.57 versus 3.0 mg.ml^−1^) and Vmax (30.8 versus 11.0 nmol of GalA.min^−1^.μg^−1^). Protein structures were determined by X-ray crystallography with the final models’ geometry, processing and refinement statistics summarized in Table 1. We solved the crystal structure of PGLR (429 aa, 1-18 and 409-429 aa not modelled, PDB: 7B7A) at a resolution of 1.3 Å using molecular replacement with 1RMG (Fig. 1*A* and SI Appendix, Fig. S2*A*) (23). PGLR crystallised as a single molecule in a P1 asymmetric unit. The crystal structure of ADPG2 (420 aa, 1-41 and 406-420 aa not modelled, PDB: 7B8B) was resolved at a resolution of 2.0 Å using PGLR as the search model (Fig. 1*A* and SI Appendix, Fig. S2*B*). ADPG2 crystals belonged to the orthorhombic space group P2_1_2_1_2_1_ with chains A and B having a Cα root mean square deviation (rmsd) of 0.924 Å. PGLR and ADPG2 fold in right-handed parallel β-helical structure, which is common to pectinases (Fig. 1A) (12). This β-helix is formed by three repeating parallel β-sheets - PB1, PB2 and PB3 which contain 11, 12, and 11 parallel β-strands respectively, as well as a small β-sheet, PB1a, having only 3 β-strands (SI Appendix, Fig. S3*A* and *B*). T1-turns, T1a-turns, T2-turns and T3-turns connect the PB1-PB2, PB1-PB1a, PB2-PB3 and PB3-PB1 β-sheets, respectively (SI Appendix, Fig. S3*C* and *D*) (24). PGLR and ADPG2 show a α-helix at the N-terminus, interacting with the T1 turn through the establishment of a disulphide bridge (PGLR, C46-C76, ADPG2, C71-C98), which shields the hydrophobic core of the enzyme (25). Superimposition of PGLR and ADPG2 structures resulted in a rmsd of 2.299 Å, predominantly due to a deviation in the region surrounding the active site, in particular N130–P142 (T3 turn, PGLR numbering) and Y304-V318 (T1a turn, PGLR numbering). Between these loops, a large cleft (10.29 Å wide for PGLR and 14.46 Å for ADPG2), open at both sides is present, exposing PB1 for accommodating the substrate and identifying PGLR and ADPG2 as putative endo-PGs (8, 12, 15).

**Table 1.** Data collection, processing and refinement statistics for PGLR and ADPG2. Statistics for the highest-resolution shell are shown in parentheses.

| Characteristic | PGLR | ADPG2 |
| --- | --- | --- |
| Data collection |  |  |
| Diffraction source | PROXIMA1A | PROXIMA1A |
| Wavelength (Å) | 0.978 | 0.978 |
| Temperature (°C) | 100.15 | 100.15 |
| Detector | PILATUS3 6M | EIGER 16M |
| Crystal to detector distance (mm) | 190.0 | 279.3 |
| Rotation range per image (°) | 0.1 | 0.1 |
| Total rotation range (°) | 360 | 360 |
| Crystal data |  |  |
| Space group | P1 | P 2<sub>1</sub>2<sub>1</sub>2<sub>1</sub> |
| a, b, c (Å) | 38.97, 41.83, 63.33 | 71.78, 88.56, 113.87 |
| α, β, γ, (°) | 93.25, 99.86, 114.95 | 90.00, 90.00, 90.00 |
| Subunits per asymmetric unit | 1 | 2 |
| Data statistics |  |  |
| Resolution range (Å) | 33.33 - 1.3 | 44.61 - 2.0 |
| Total No. of reflection | 761821 (55201) | 645204 (65696) |
| No. of unique reflection | 83668 (8043) | 47381 (4654) |
| No. of reflections, test set | 4182 (401) | 2368 (233) |
| R<sub>merge</sub> (%) | 7.64 (77.7) | 8.9 (97) |
| Completeness (%) | 96.1 (92.0) | 99.9 (99.3) |
| (I/σ(I)) | 16.24 (2.78) | 16.91 (2.56) |
| Multiplicity | 9.1(6.9) | 13.6 (14.0) |
| CC<sub>1/2</sub> (%) | 99 (86.3) | 99 (92.1) |
| Refinement |  |  |
| R<sub>crys</sub>/R<sub>free</sub> (%) | 14.2 / 17.7 | 18.9 / 23.0 |
| Average B - factor (Å<sup>2</sup>) | 29.1 | 27.89 |
| No. of non-H atoms |  |  |
| Protein | 3085 | 5563 |
| Ion | - | 5 |
| Ligand | 100 | - |
| Water | 609 | 999 |
| Total | 3794 | 6567 |
| R.m.s. deviations |  |  |
| Bonds (Å) | 0.015 | 0.006 |
| Angles (°) | 1.59 | 1.06 |
| Ramachandran plot |  |  |
| Most favoured (%) | 94.6 | 93.58 |
| Allowed (%) | 5.4 | 6.28 |
| Outlier (%) | - | 0.14 |

**Figure 1.**
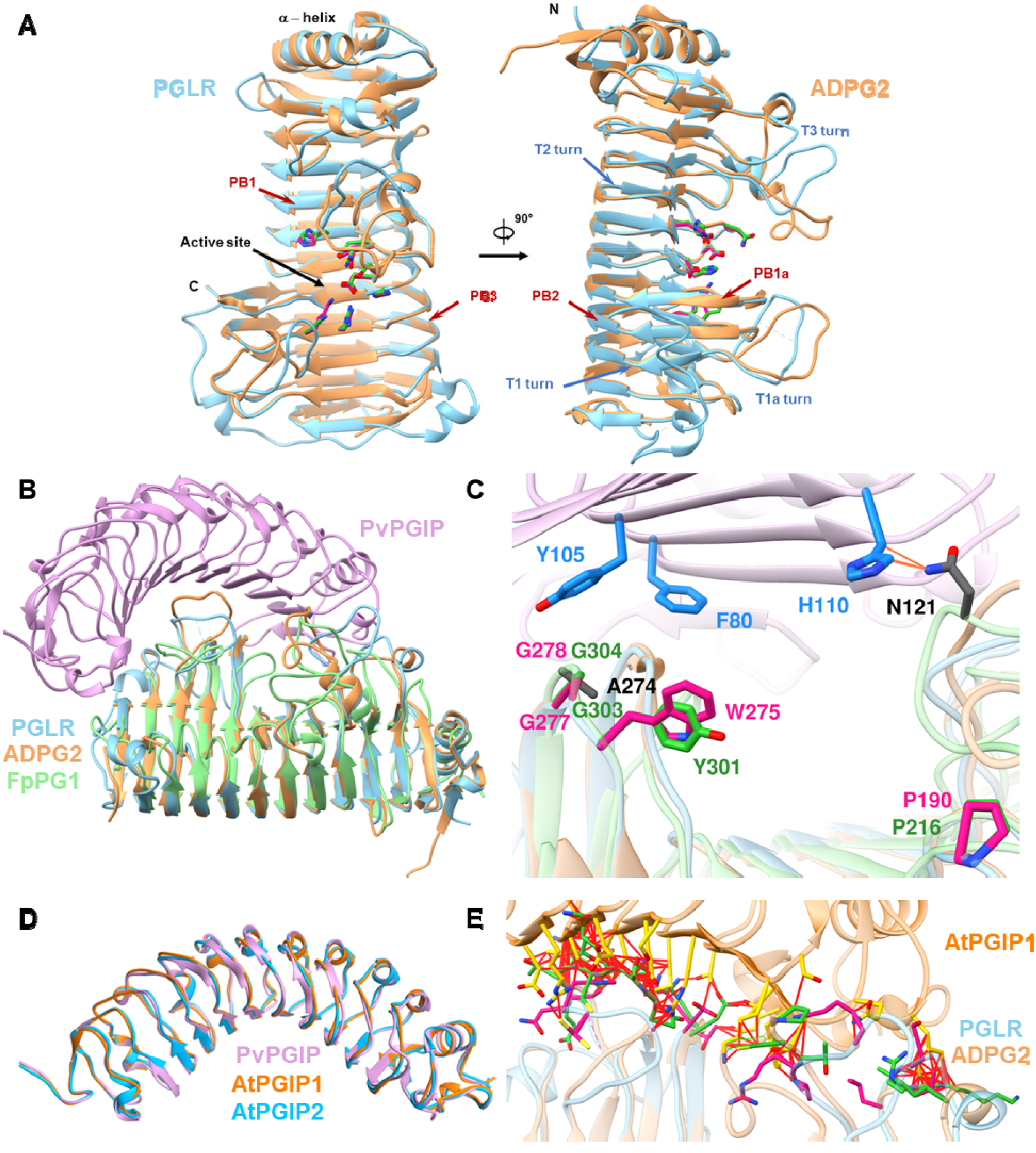
Structure comparison of PGLR and ADPG2 and identification of novel amino-acids required for activity. A) Overall structure of PGLR and ADPG2 represented in ribbon diagrams which are colored in blue and brown, respectively. Right-handed parallel β-helical structure consisting of β-strands (red) and turns (blue). PGLR and ADPG2 active site amino acids are pink and green colored. β-sheets are turns are indicated by red and blue arrows. B) Ribbon representation of *Phaseolus vulgaris* PGIP2 (PvPGIP2, plum), PGLR (blue), ADPG2 (brown), *Fusarium phyllophilum* PG (FpPG1, green). C) Detailed representation of aa involved in PvPGIP2-FpPG1 interaction (PvPGIP2 amino-acids in blue and FpPG1 amino-acids in grey), with orange lines representing van der Waals contacts. Key aa (N121, A274) mediating the interaction in FpPG1 are absent in PGLR and ADPG2. D) Superimposition of crystallised PvPGIP2 with models of Arabidopsis PGIP1 (AtPGIP1, orange) and PGIP2 (AtPGIP2, blue). E) Interactions of AtPGIP1 with PGLR and ADPG2. Amino acids of AtPGIP1 (yellow), PGLR (pink) and ADPG2 (green) included in clashes closer than 0.6 Å are shown. The red lines represent atoms overlap of minimum 0.6 Å

### Structural determinants of absence of plant PG-plant PGIP interactions

While PGLR and ADPG2 show low sequence identity with fungal enzymes (sequence identity: 19%-25% with *Aspergillus aculeatus* (AaPG1), *Aspergillus niger* (AnPG1 and AnPGII), *Fusarium phyllophilum* (FpPG1), *Pectobacterium carotovorum* (PcPG1), *Chondrostereum purpureum* (CpPG1)), they show high structural similarity with a rmsd of 4.753 to 7.761 Å between all atoms (SI Appendix, Fig. S4*A* and *B*). Still, PGLR does not interact with plant PGIPs, as shown by the lack of inhibition of PGLR activity by *Phaseolus vulgaris* PGIP2 (PvPGIP2), while this interaction exists with fungal PGs (8, 18). To understand the structural basis of this absence of inhibition of plant PG activity by PGIP we superimposed the resolved structures of PGLR and ADPG2 onto the *Fusarium phyllophilum* PG (FpPG1) - PvPGIP2 complex (Fig. 1*B*) (16, 18). In FpPG1, a S120-N121-S122-N123 stretch, within the protein’s N-terminal loop, plays a key role in the PG-PGIP interaction (N121 notably interacting with H110 of PvPGIP2). PGLR and ADPG2 N-terminal loops are, on the other hand, rich in bulkier and chemically different residues, including M132, M133 and M137 for PGLR and K160, K162 and K166 for ADPG2 (SI Appendix, Fig. S5). At the C-terminus, A274, the aa that contributes to hydrophobic-stabilizing interactions for the FpPG1-PvPGIP is replaced by G277/G278 and G303/G304 in PGLR and ADPG2, respectively (Fig. 1*C*) (18). Moreover, plant PGs have a specific H to P (P190/P216) substitution together with W275/Y301 insertion which can hinder the PG-PGIP interaction (26). We next modelled AtPGIP1 and AtPGIP2, which superimpose to PvPGIP2 with a rmsd of 1.194 and 1.201 Å, respectively (Fig. 1*D*). The analysis of the models for PGLR/ADPG2-AtPGIP1/AtPGIP2 complexes showed that multiple aa are involved in steric clashes (between 81 and 275 atom contacts depending of the PG-PGIP pair, SI Appendix, File. S1), which, together with the above-mentioned structural features, can explain the absence of the interaction between PGs and PGIPs from Arabidopsis, and lack of protein-mediated inhibition of PG activity in planta (Fig. 1*E* and SI Appendix, Fig. S6*A* and *B*).

### PGs with conserved active sites show differences alongside the binding groove subsites known to be of importance for substrate interaction and processivity

Comparison of PGLR and ADPG2 sequences and structures with those from bacteria and fungi reveal that the active site is formed by four conserved structural motifs NTD, DD, GHG, RIK positioned at subsites -1 and +1 of the PB1 (27–29). Eight of these aa N191/N217, D193/D219, H196/H222 D214/D240, D215/D241, H237/H263, R271/R297, K273/K299 (PGLR/ADPG2 numbering) are strictly conserved with the three aspartates responsible for the hydrolysis of the substrate (Fig. 2*A*) (11, 25, 28, 30). To determine the importance of specific aa, five site-directed mutations were designed for PGLR: D215A occurring in the active site, R271Q (subsite +1), and the histidine mutants H196K, (subsite -1), H237K (subsite +1) and H196K/H237K (SI Appendix, Fig. S7). Histidine residues could potentially modulate the activity of the enzyme by controlling the protonation state of residues placed in subsites flanking the hydrolysis site (Fig. 2*A*). Their activities, on PGA, and dissociation binding constants (Kd) to the substrate (represented by a mix of OGs of mean DP12 and DM5 on which PGLR shows activity, SI Appendix, Fig. S8) were determined by MicroScale Thermophoresis (MST). D215A and R271Q mutations resulted in a total loss of activity with a substantial reduction in binding affinity (Kd of 2567 nM and 4840 nM for D215A and R271Q, respectively compared to 1246 nM for WT, Fig. 2*B*). While binding affinities of all histidine mutants were not significantly different, the H237K and H196K/H237K mutants showed no residual activity while the H196K mutant had featured only 22% residual activity of the WT. Although having conserved active sites, sequence and structure analyses showed that twelve aa positioned alongside the binding groove (subsites from -5 to +5), previously shown to be of importance for substrate interaction and processivity differ between PGLR and ADPG2 (Fig. 2C-D) but as well with those of the fungal AaPG1 (SI Appendix, Fig. S9) (12, 27, 29). For instance, at subsite -5, PGLR harbours R146, that can be responsible for the interaction with a carboxylate group of GalA, while ADPG2 harbours T172. Similarly, at subsite -4, Q198 in PGLR is replaced by T224 in ADPG2. At subsite -4, -3 and -2 a patch formed by Q198, Q220 and the positively charged K246 in PGLR is mutated into T224, E246 and D272 in ADPG2. At subsite -1 S269 in ADPG2, that can form hydrogen bonds with the substrate is mutated into G243 in PGLR. Finally, at subsite +2 and +3 D293 and K322 in ADPG2 are replaced by T267 and A296 in PGLR.

**Figure 2:**
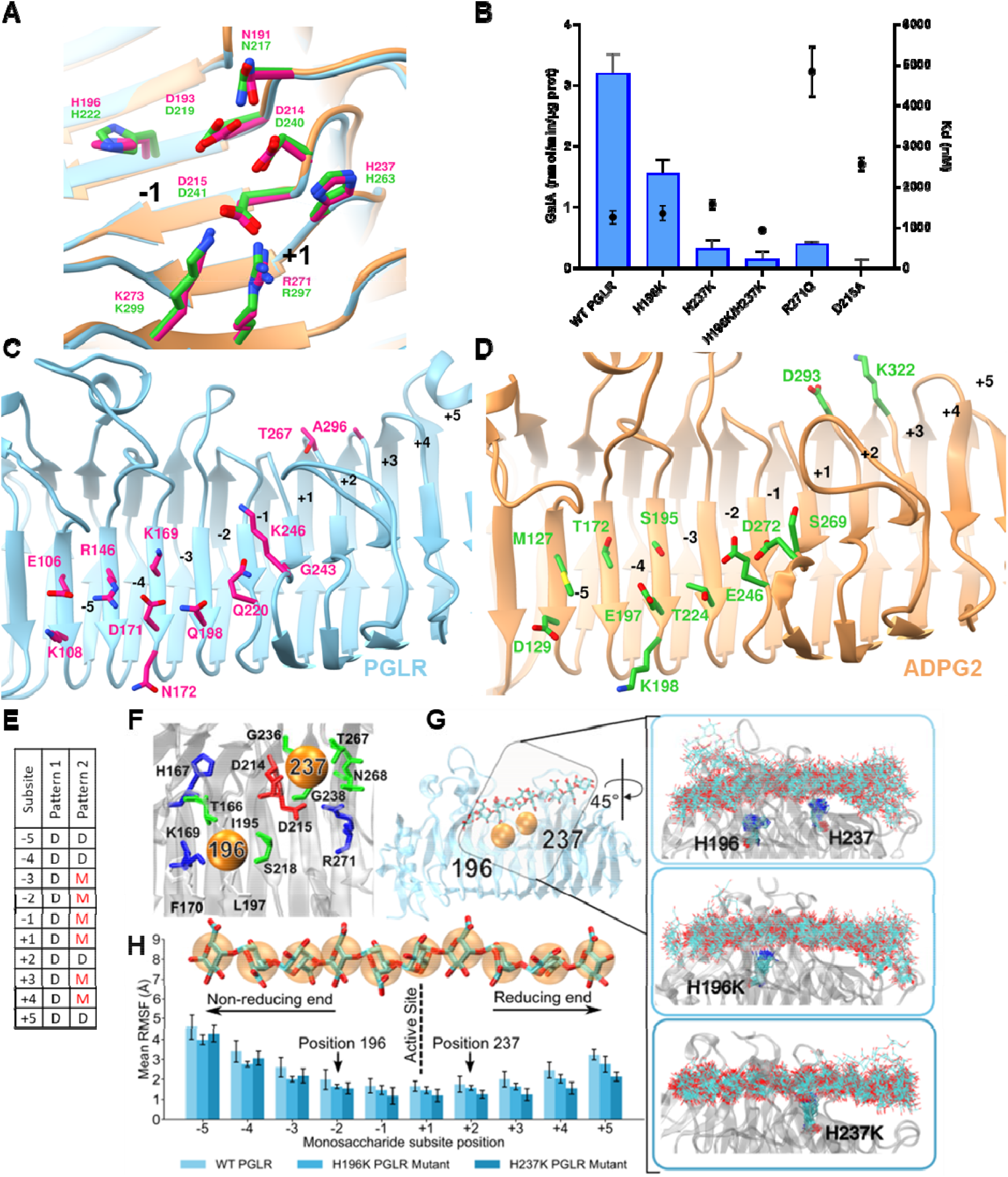
Structure of the PGLR-ADPG2 active site and binding groove. Role of H196 and 237 for PG activity. A) Active site of the PGLR and ADPG2 highlighting absolutely conserved aa. D193, D214 and D215 are aa involved in substrate hydrolysis. Black numbers indicate the subsites. B) Total PG activity of WT and mutated forms of PGLR (H196K, H237K, H196K/H237K, R271Q, D217A) on PGA (blue bars) and MST analysis of the interaction between WT and mutated forms of PGLR using a substrate of DP12 and DM5 (black dots). C) Structure of PGLR binding groove (subsite -5 to +5). D) Structure of ADPG2 binding grove (subsites -5 to +5). E) Sequence of the fully de-methylesterified (pattern 1) or 60% methylesterified (pattern 2) decasaccharides simulated in complex with ADPG2 and PGLR. D: de-methylesterified GalA, M: methylesterified GalA. F) cross-section of the substrate binding groove highlighting the positions of H196 and H237, which are represented as orange spheres. Positively and negatively charged residues are shown in blue and red, respectively, while polar residues are shown in green and represented as sticks. G) PGLR in complex with a decasaccharides substrate, in the insets the conformational ensembles of the substrate and of H196, H237 and H196/H237 are shown, by reporting conformations obtained every 10 ns. H) Root mean square fluctuations (RMSF) of each monosaccharide across the binding groove for WT PGLR and its mutants.

### Molecular dynamic (MD) simulations reveal distinct substrate-dependent dynamics of PGLR and ADPG2

The large number and chemical diversity of interactions across the binding groove make structural comparisons between different PG isoforms poorly informative. Such a diversity can result in different dynamic behaviours of enzymes and/or substrates, which could translate into different functional profiles. We performed MD simulations on PGLR and ADPG2 in complex with either a fully de-methylesterified (pattern 1) or 60% methylesterified (pattern 2) decasaccharides (Fig. 2*E*), able to occupy the totality of the binding groove (subsites from -5 to +5). We first simulated PGLR, as well as H196K and H237K mutants in complex with fully de-methylesterified decasaccharides, and the analysis of substrate dynamics, through the quantification of subsite-specific root mean square fluctuations (RMSF), revealed a trend between enzymatic activity (Fig. 2*B*), substrate dynamics (Fig. 2*F*-*H*) and the total number of contacts between the substrate and enzymes (SI Appendix, Fig. S10). MD simulations of the PGLR mutants (H196K and H237K) revealed how substrate dynamics is affected all along the binding groove, even with a single histidine mutation occurring in subsites either towards the non-reducing end (H196K – subsite -1) or the reducing end of the sugar (H237K – subsite +1). Overall, a rigidification of the substrate coincides with the loss of activity observed in experiments (Fig. 2*B*), with the H237K mutant (total loss of activity) showing the lowest RMSF in subsites -1 to +5 compared to the H196K (22% residual activity) and the WT (highest substrate dynamics, Fig. 2*G* and *H*).

The substrate dynamics can be also seen when comparing the RMSF of ADPG2 and PGLR when in complex with either de-methylesterified or methylesterified decasaccharides. For both enzymes, de-methylesterified oligomers are overall less dynamic, hence more tightly bound in the binding groove (Fig. 3*A* and *B*). Quantitative differences in the RMSF of the two complexes suggest that, for the same substrate either being de-methylesterified or partially methylesterified, the binding to PGLR is tighter. Moreover, for each substrate, ADPG2 retains a higher activity compared to PGLR (SI Appendix, Fig. S1*E*), which again corroborates the observation that methylesterified substrates are overall less dynamic in complex with PGLR when compared to ADPG2 (Fig. 3*A* and *B*). The observed substrate dynamics is linked to the total number of contacts with the enzyme, with some noticeable differences between the two isoforms. When in complex with de-methylesterified substrates, both enzymes establish a larger number of contacts with the oligosaccharides. PGLR has however the ability to make a larger number of contacts, which is especially relevant for salt-bridges and hydrogen-bonds (Fig. 3*C*). The reduced substrate dynamics when bound to histidine PGLR mutants is also reverberated into a higher number of contacts (SI Appendix, Fig. S10). A comparison of the enzymatic motions revealed that PGLR and ADPG2, while engaged to the same decasaccharide substrate, explore separate conformational states, which are especially related to the fluctuations of unstructured regions flanking the binding groove. While for PGLR these are the regions flanking the substrate’s non-reducing end (residues K108, R146, K169), in the case of ADPG2 they flank the binding cleft and in proximity of the substrate’s reducing end (SI Appendix, Fig. S11*A* and *B*). Relevant differences can also be observed between the electrostatic potentials of the two enzymes, calculated by solving the Poisson-Boltzmann equation in implicit solvent (SI Appendix, Fig. S11*C* and *D*). Compared to ADPG2,

**Figure 3.**
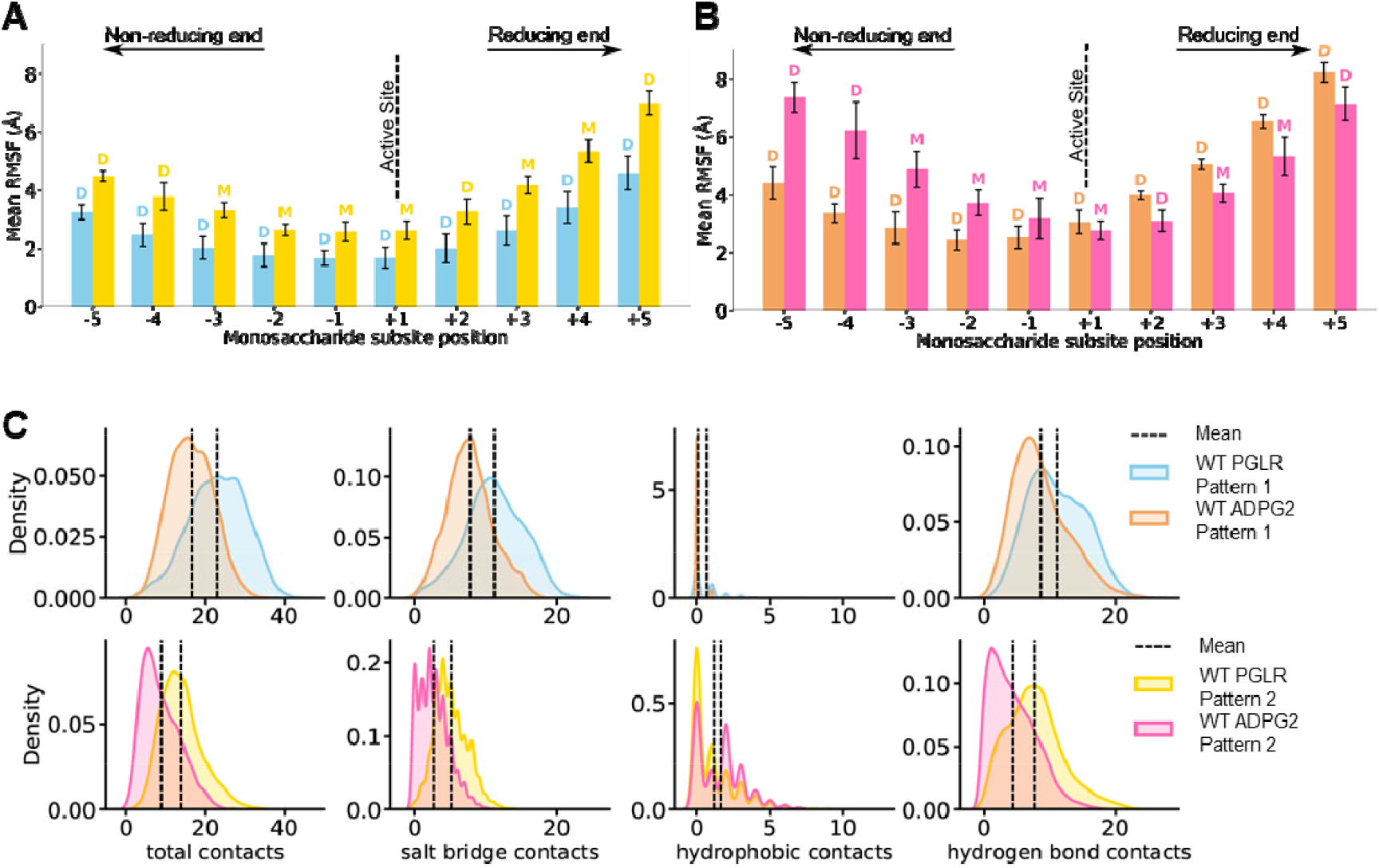
PGLR and ADPG2 show distinct substrate dynamics. A-B) Root mean square fluctuations (RMSF) of each monosaccharide bound across the binding groove of PGLR (A) or ADPG2 (B). In each panel, fully de-methylesterified (pattern 1 – cyan in A and orange in B) or 60% methylesterified decasaccharides (pattern 2 – yellow in A and pink in B) are shown. C) Analysis of the contacts between PGLR or ADPG2 and substrates either fully de-methylesterified (pattern 1) or characterized by 60% methylesterification (pattern 2).

PGLR shows a much more positively charged electrostatic potential within the substrate binding cleft, in line with pronouncedly reduced dynamics for a negatively charged (de-methylesterified) substrate, which would undergo much stronger electrostatically dominated interactions with the enzyme. Overall, subtle differences within the amino acid composition of certain subsites can convey specifically different activity profiles from a seemingly identical fold, which is likely to generate distinct substrate binding affinities, and end-products.

### The differences in PGLR and ADPG2 binding’s kinetics leads to specific pools of pectin-derived fragments

The calculated RMSF shows differences in enzyme-substrate dynamics once the substrate is bound, which could reflect differences in the binding affinities of the enzymes towards specific substrates. Using fluorescence-based switchSENSE® aptasensor, we determined binding kinetics for enzyme-substrate interactions for both PGLR and ADPG2, by quantifying substrate association (k_on_) and dissociation (k_off_) rate constants, as well as equilibrium dissociation constant (K_D_) using substrates with various DPs and DMs (PGA, pectins DM 20-34%, OGs of DP12DM5, DP12DM30, DP12DM60, Table 2). ADPG2 displayed affinities much higher for low-DM substrates (i.e. PGA and DP12DM5) than those determined with the high-DM pectins (K_D_ *ca*. 10 to 60 times lower; Table 2) and comparable to those of PGLR. Considering the kinetics constants, PGLR and ADPG2 show no difference for k_on_ for pectins of low DM, including PGA and DP12DM5 (1320/1120 and 953/833 M^−1^s^−1^), respectively. In contrast, when the DM of the substrate is increased (DM 20-34%, DP12DM30 and DP12DM60), the k_on_ is always higher (∼x 3 to 16 times) for PGLR compared to ADPG2. This suggest that for methylesterified pectins, PGLR, in line with the MD simulations and lower RMSF compared to ADPG2, associates much tighter with the substrate. This is as well reflected by the lower K_D_ determined for PGLR compared to ADPG2. No such drastic differences are measured for k_off,_ as values for PGLR and ADPG2 are in the same range for most substrates.

**Table 2.**
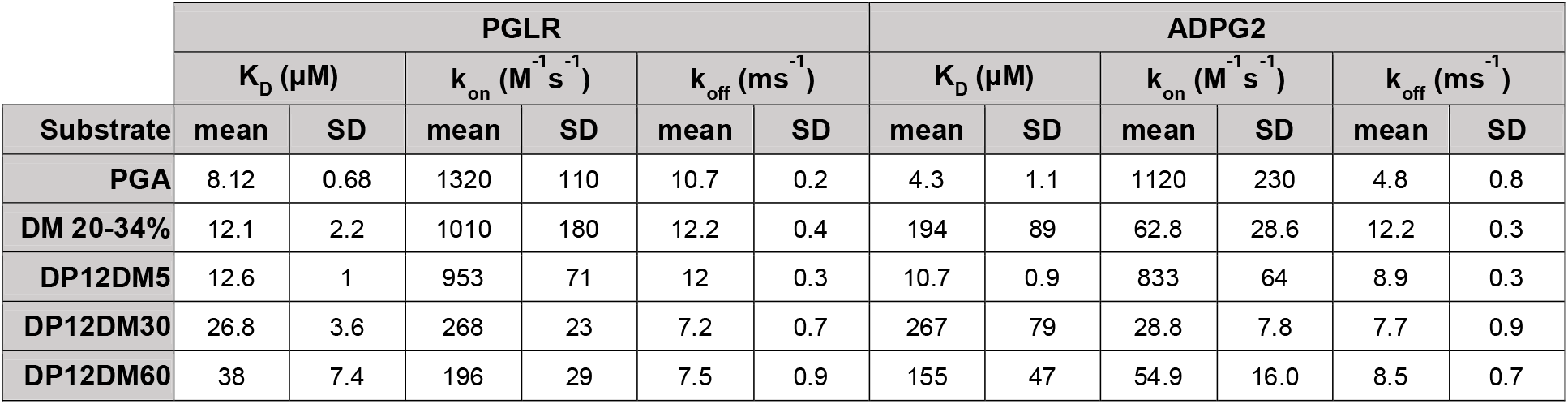
kon, koff and kD measurements for PGLR and ADPG using substrates of various degrees of polymerization. PGA: Polygalacturonic acid, pectins DM 20-34%: Commercial pectins of DM30%, DP12DM5/DP12DM30/DP12DM60: Pool of OG cantered on DP12 with increasing DM (5%, 30%, 60%). Data shown average of three replicates.

| Substrate | PGLR |  |  |  |  |  | ADPG2 |  |  |  |  |  |
| --- | --- | --- | --- | --- | --- | --- | --- | --- | --- | --- | --- | --- |
| | $K_D$ ($\mu M$) | | $k_{on}$ ($M^{-1} s^{-1}$) | | $k_{off}$ ($ms^{-1}$) | | $K_D$ ($\mu M$) | | $k_{on}$ ($M^{-1} s^{-1}$) | | $k_{off}$ ($ms^{-1}$) | |
|  | mean | SD | mean | SD | mean | SD | mean | SD | mean | SD | mean | SD |
| PGA | 8.12 | 0.68 | 1320 | 110 | 10.7 | 0.2 | 4.3 | 1.1 | 1120 | 230 | 4.8 | 0.8 |
| DM 20-34% | 12.1 | 2.2 | 1010 | 180 | 12.2 | 0.4 | 194 | 89 | 62.8 | 28.6 | 12.2 | 0.3 |
| DP12DM5 | 12.6 | 1 | 953 | 71 | 12 | 0.3 | 10.7 | 0.9 | 833 | 64 | 8.9 | 0.3 |
| DP12DM30 | 26.8 | 3.6 | 268 | 23 | 7.2 | 0.7 | 267 | 79 | 28.8 | 7.8 | 7.7 | 0.9 |
| DP12DM60 | 38 | 7.4 | 196 | 29 | 7.5 | 0.9 | 155 | 47 | 54.9 | 16.0 | 8.5 | 0.7 |

To determine whether the differences in subsites structure, enzyme dynamics and binding affinities can translate into differences in the processivity of PGLR and ADPG2, we assessed the products generated by either of the enzymes. Using PGA as a substrate, PGLR or ADPG2 maximum activities were reached after 1-hour digestion, generating products that cannot be further hydrolysed. ADPG2 total activity was higher than that measured for PGLR. Furthermore, the addition of ADPG2 following a first hour substrate incubation with PGLR, led to an increase in total PG activity, confirming putative differences in processivity between the two enzymes, ADPG2 being able to hydrolyse PGLR’s end-products (Fig. 4*A*). We then used a recently developed LC-MS/MS oligoprofiling approach (31) to analyse the reaction products and confirmed, using PGA as a substrate, that both enzymes have endo activities, as suggested by the structural features of the binding cleft, and that ADPG2 releases higher proportion of short-sized OGs (<DP4) compared to PGLR (Fig. 4*B*). On pectic substrates of DM 20-34%, the pool of OGs produced by PGLR differed to that of ADPG2 (Fig. 4*C*). In particular, PGLR released de-methylesterified OGs of DP5 to DP9, as well as specifically methylesterified forms of more than 6 GalA units that were either poorly represented or absent in the pool of end-products produced by ADPG2. The main products of ADPG2 were indeed de-methylesterified OGs of DP2 to DP4, as well as large amount of GalA4Me (Fig. 4*C* and Fig. 4*C*, Inset). When comparing the OGs produced by PGLR, ADPG2 and AaPG1 upon enzymatic activity on pectins with DM between 20 and 34% using principal component analysis (PCA), PGLR and AaPG1 were separated according to the first dimension (Dim1 54.6.7% of the variance) while ADPG2 clustered according to second dimension (Dim2 40.4% of the variance), with main loadings being, as an example, GalA2, GalA3, GalA4Me2, GalA9Me (SI Appendix, Fig. S12*A* and *B*). Overall, ADPG2 and PGLR have nearly identical folds that, through distinct subsite structure and enzymes’ dynamics, could translate into different enzymatic processivities. Indeed, PGLR and ADPG2 differ in their intrinsic processivities, P^Intr^, being described as the average number of consecutive catalytic acts before enzyme-substrate dissociation. P^Intr^ is dependent on the dissociation probability, P_d_, calculated using the turnover number (k_cat_) and rate constant of dissociation (k_off_) (32). P_d_ values were 4.8 × 10^−4^ and 5.1 × 10^−5^, and P^Intr^ values were 2081 and 19777, for PGLR and ADPG, respectively. This data shows that, albeit acting both as processive enzymes (P_d_<<1), PGLR and ADPG2 differ in the extent by which they act on the substrate, with ADPG2 being much more processive than PGLR, as reflected by the lower size of the released products detected with LC-MS/MS.

**Figure 4.**
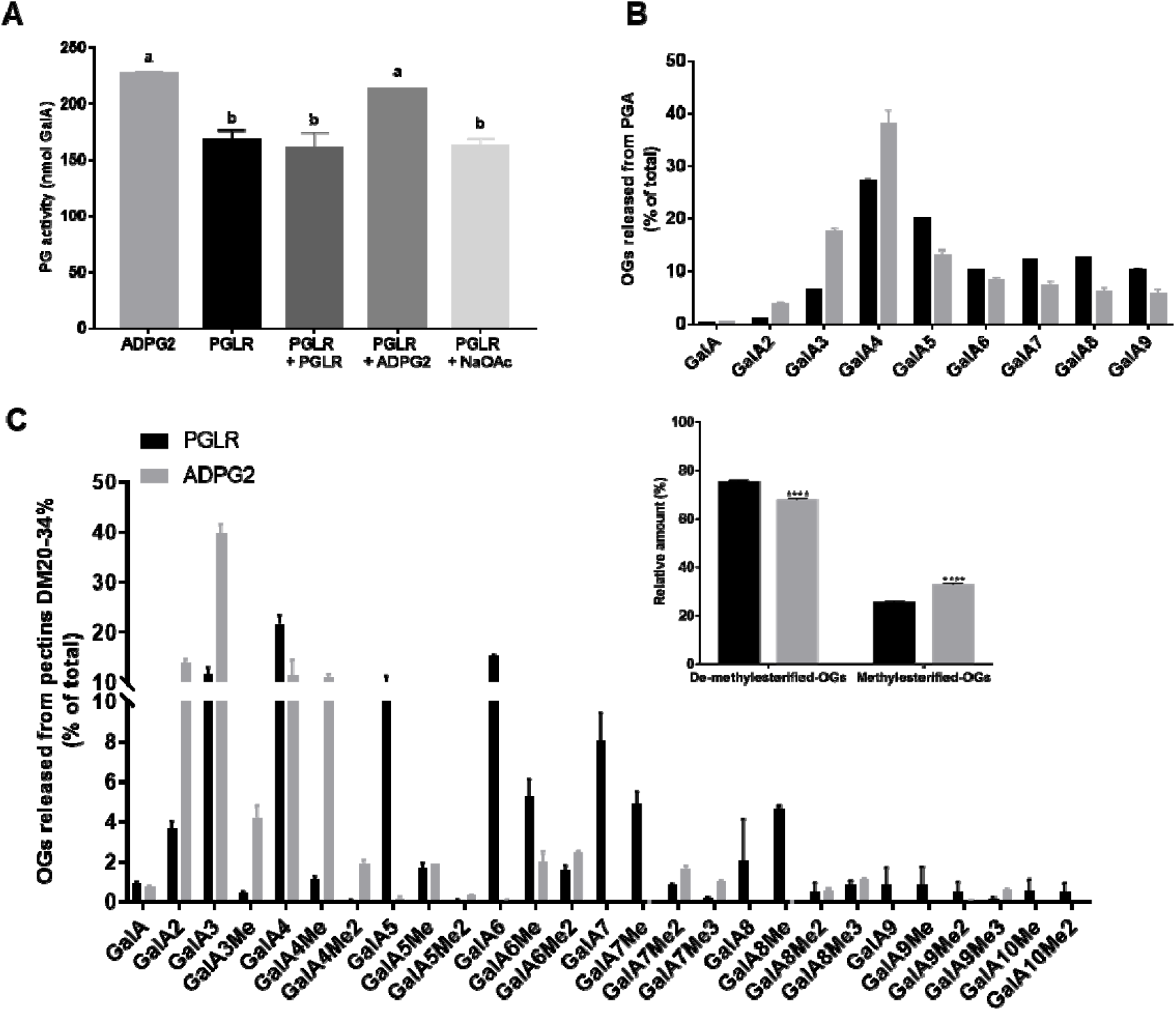
PGLR and ADPG2 release distinct OGs. (A) Activity tests performed on PGA (DM 0) after 1-hour digestion by ADPG2, PGLR and by adding PGLR or ADPG2 for 1 hour after a first digestion by PGLR. NaOAc (sodium acetate): negative control. (B) Oligoprofiling of OGs released after 1-hour digestion of PGA by PGLR (black) or ADPG2 (grey) at 40°C, pH 5.2. C) Oligoprofiling of OGs after overnight digestion of pectins DM 20-34% by PGLR (black) or ADPG2 (grey) at 40°C, pH 5.2. Inset: Cumulative OGs released by PGLR and ADPG2 after over-night digestion on pectins DM 20-34% at 40°C, pH 5.2. Two-way ANOVA with Sidask’s multiple comparison test, P value ****<0.0001.

### The exogenous application of PGs with different processivities translate into distinct effects on root development

Considering the localization of the expression of *PGLR* and *ADPG2* during root development, we tested the activity of both enzymes on root cell walls, whose pectins can be both methylesterified and acetylated (21, 22, 33). Noticeably, PGLR released a higher proportion of acetylated OGs (including GalA5Ac, GalA6Ac, GalA6Ac2) compared to ADPG2, in addition to longer oligomers on average (Fig. 5*A* and Fig. 5*A*, Inset). Similarly, to what was observed on methylesterified pectins, the main OGs produced by ADPG2 were of lower DPs as compared to what produced by PGLR, corresponding mainly to unsubstituted GalA2 and GalA3. As a read-out, and to determine how distinct processivities of PGLR and ADPG2 on HG can translate into distinct phenotypes *in muro*, we assessed the effects of exogenously-applied purified enzymes on developing roots. Iso-activities of PGLR and ADPG2 were added in the culture medium of 6-day-old seedlings, for either one or three days, and phenotypical changes were examined. If one day’s application of either of the enzymes did not affect root length, ADPG2 significantly impaired root elongation when applied for three days (Fig. 5*B*). In contrast, in the latter condition, a slight effect was measured for PGLR albeit non-significant. (Fig. 5*B*). To determine more precisely if cells are differentially affected upon enzyme application depending on their spatial positioning (meristematic, elongating or fully elongated), we then measured the length of the firsts 50 cells from the root tip after three days of enzymes’ application, using EGFP-LTI6b reporter line that specifically labels plasma membrane (Fig. 5*C*) (34). Cell length was not affected by the application of either of the enzymes up to the 40^th^ cell. In contrast, the application of ADPG2 drastically reduced the length of the cells in the elongation zone as early as cell 40; while the effects measured for PGLR were from cell 46 onwards and were lower compared to that of ADPG2 (Fig. 5*D*). Further differences between the enzymes can be highlighted by analysing their effects on the morphology of the root cap, the structure at the tip of the root which supports growth and protects the root meristem. The application of ADPG2 for three days had much drastic effects on root cap detachment as compared to that of PGLR suggesting that it has more drastic effects on cell-to cell adhesion (Fig. 5*E*). Altogether, this shows that the biochemical specificities/processivities of the two enzymes will ultimately translate into distinct effects on development, when applied exogenously in the culture media.

**Figure 5.**
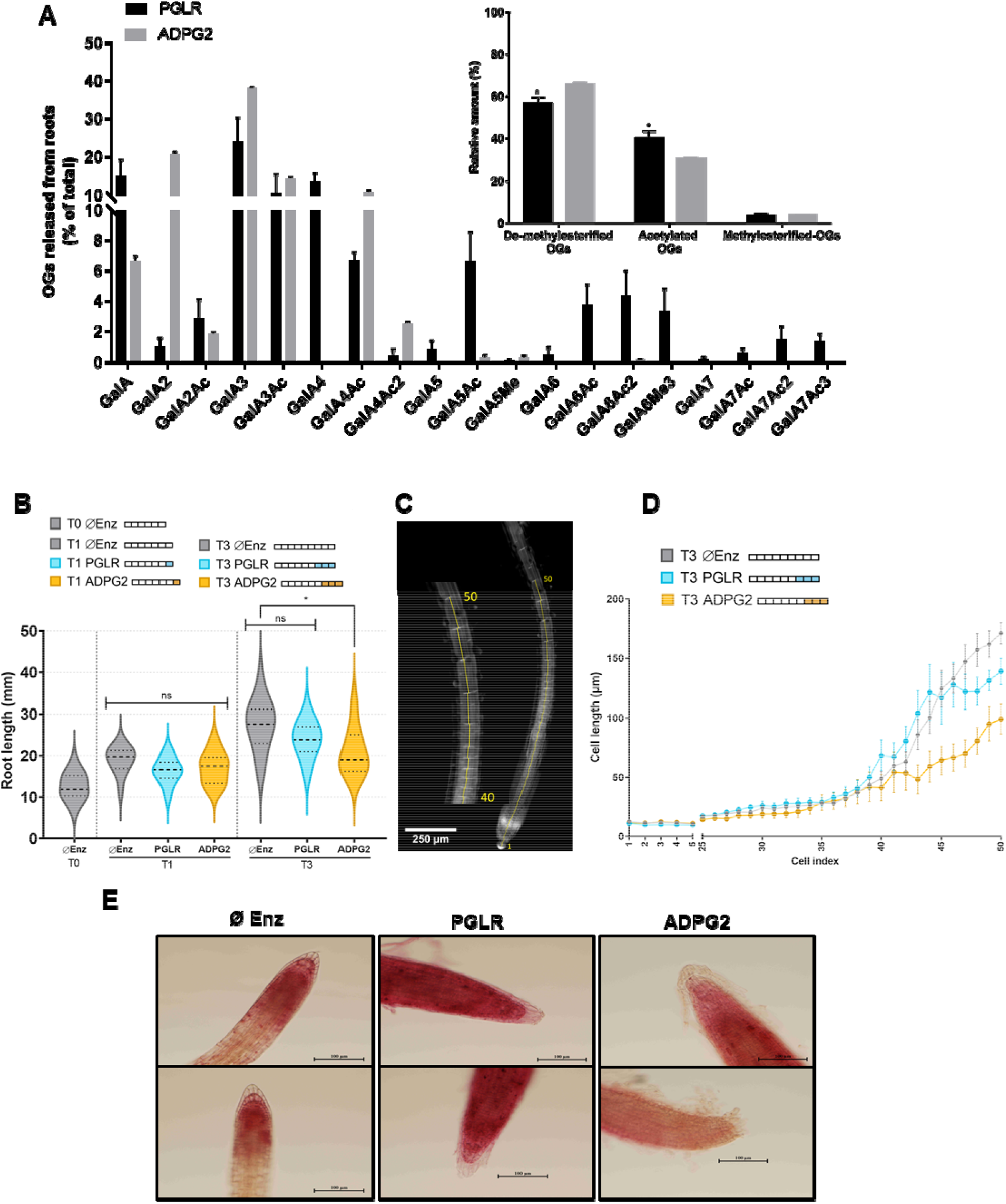
PGLR and ADPG2 are active on roots pectins and have distinct effects of root length. A) Oligoprofiling of OGs after digestion of roots cell wall by PGLR (black) and ADPG2 (grey) at 40°C, pH 5.2 after over-night digestion, (Inset: Cumulative OGs released by PGLR (black) and ADPG2 (grey) after over-night digestion of roots cell walls at 40°C, pH 5.2). Two-way ANOVA with Sidask’s multiple comparison test, P value *0.0290. B) Effects of the exogenous application of PGLR and ADPG2 on total root length of Arabidopsis seedlings. PGLR and ADPG2 were applied at iso-activities for one or three days on 6-day-old seedlings grown in liquid media. C) Root cell numbering using EGFP-LTI6b reporter lines. D) Effects of 3-day exogenous application of PGLR and ADPG2 on the cell length of the first 50 roots cells of 7-day-old seedlings. E) Effects of 3-day exogenous application of PGLR and ADPG2 on root cap structure of 7-day-old seedlings (2 representative images per condition). Buffer (Ø Enz) was used as negative control. Scale bar represents 100 mm.

## Discussion

PGs play an important role in the control of pectin chemistry, contributing to changes in the cell wall mechanics, with important consequences on plant development (4–7). In Arabidopsis, PGs are encoded by 68 genes: an abundance which is hard to rationalize within the context of the plant cell wall. Here we elucidated the structure-to-function relationships for two plant PGs, PGLR and ADPG2, whose expression overlaps in Arabidopsis roots. Both enzymes have nearly identical triple β-helix folds commonly found in other pectinases, including fungal endo-PGs (12, 25, 28), pectin/pectate lyases (24, 35, 36) and rhamnogalacturonases (23), with a large cleft opened at both sides that accommodates oligomeric substrates and confirms that PGLR and ADPG2 are endo-PGs (25). The resolution of the crystal structure for plant PGs first rationalized the structural determinants of the absence of inhibition of plant enzymes by plant PGIPs, as PGLR activity was indeed not inhibited by *P. vulgaris* PGIP2 (PvPGIP2) (8). Structurally, the key aa of *F. phyllophilum* FpPG1 (S120-N121-S122-N123) needed for determining the interaction of this pathogenic PG with PvPGIP2, are absent in the T3 loop of PGLR and ADPG2. The homology modelling of Arabidopsis AtPGIP1 and AtPGIP2 further highlighted the absence of PGIP-mediated regulation of endogenous PG activity in plants as, albeit having highly conserved structure with that of PvPGIP2, they are lacking H110 and Q224 residues, required for inhibition (37). This suggests that cellular regulation of PG is mediated by other means at the cell wall, one of which being, as demonstrated in this study, the differential processivities of the enzymes.

The main challenge in understanding subtle differences between isoforms of PGs and other carbohydrate binding enzymes (CBEs) are mostly related to the large binding interface that characterizes the interaction between CBEs and oligomeric substrates. We tackled this challenge by designing strategic mutations across the binding cleft of the structurally characterized PGLR and functionally analysing the enzymes with combined computational and experimental methodologies. Our findings confirmed the importance of D215 for substrate hydrolysis, as well as R271 in binding and positioning the substrate at the catalytic subsite +1, as previously reported for fungal PGs (25, 30). Besides residues actively important in stabilizing the substrate, we find that other interactions in subsites flanking the catalytic subsite, crucially regulate substrate dynamics and correlate with enzymatic activity. Histidine to lysine mutants in PGLR (H196K, H237K and H196K/H237K), that might generally be important in controlling the observed pH-dependent activity of other PGs, show how the distribution of charges affects substrate dynamics. Most interestingly, substrate rigidification reported by MD upon the insertion of a positive charges, increases the number of contacts with the substrate across the substrate binding interface, negatively impacts enzymatic activity as reported by the experimental biochemical characterization of the mutants. The importance of substrate dynamics in the activity of other CBEs has been also previously reported and it might be a key factor in regulating the processive activity of CBEs more generally, with processivity being limited by substrate dissociation (38, 39).

We next investigated whether the processivities of PGLR and ADPG2 differ, which could be related to their different subsite’s composition affecting enzymes’ dynamics. For instance, D293 and K322 in ADPG2 are replaced by T267 and A296 in PGLR, which could modify the enzyme-substrate interaction and the enzyme specificity. The determination of the dynamics, measured as the RMSF, of the enzymes in complex with a decasaccharide of GalA showed that i) for a given enzyme, the enzyme’s dynamics differs with the DM of the substrate and ii) ADPG2 was overall more dynamic, with a higher RMSF, as compared to PGLR. Together with these simulations, the determination of the binding kinetics of the enzyme-substrate interactions led to hypothesizing distinct processivities for the two enzymes. When considering pectins of high DM (DM 20-34%, DP12DM30, DP12DM60), the affinities of both enzymes are k_on_-dominated, with PGLR associating much tighter with the substrate. Interestingly, the affinity of ADPG2 for the low-DM substrates is higher than that towards the high-DM pectins and is comparable to the affinities determined for PGLR. Considering the lubricating hypothesis, inferred from the studies on pectin methylesterases, and intrinsic processivity calculations, ADPG2 acts more processively on the HG chain than PGLR, and that would occur more favourably with low-DM substrates (Fig 6*A* and *B*) (35, 38). Altogether, despite overall higher Km value, these results are in accordance with the lower RMSF, the substrate being more tightly bound inside the active site, and k_on_, substrate association rate constant for the active site measured for PGLR. This would impair the sliding of the enzyme onto the chain, leading to enzyme-substrate dissociation and reiteration of enzyme attack onto the chain (Fig. 6*A*). Such distinct processivities effectively translated into different end-products, with ADPG2 releasing OGs of short DP (methylesterified or not) from either commercial substrates or root cell wall extracts, while PGLR released a high proportion of non-methylesterified OGs of higher DP (Fig. 6*B*). As highlighted by the fact that ADPG2 can hydrolyse PGLR-generated OGs, one could envisage a cooperative action of both enzymes in the cell wall to finely-tune HG structure during root development. A number of studies previously showed the impact of the changes in PG activity, either through the study of loss-of-function mutants or over-expressing lines in Arabidopsis, on developmental processes as diverse as dark-grown hypocotyl development, stomata formation and root development (3–8). However, considering the size of the *PG* gene family, functional genomics approaches (mutants and/or overexpressing lines) can lead to counter intuitive results where changes in expression of one given *PG* gene leads to either increase, or decrease of total PG activity, owing to compensation mechanisms among the gene family (8). By using exogenous application of purified enzymes, one could expect more direct effects, which can allows linking the enzymes’ processivities to their impact on pectins’ structure, on their interaction with other cell wall components, on cell wall integrity, and ultimately on plant development. The consequences of the exogenous application of the highly processive ADPG2 had indeed different effects on root development as compared to that observed with PGLR; HG remodelling upon ADPG2 application leads to strong defects in root elongation and in cell adhesion at the root cap. The root cap phenotype of ADPG2-treated roots is similar to that reported for *ROOT CAP PG1* (*RCPG1*) overexpressing lines, known to be involved in root cap removal, suggesting that enzymes might share common biochemical specificities and/or processivities (40).

**Figure 6.**
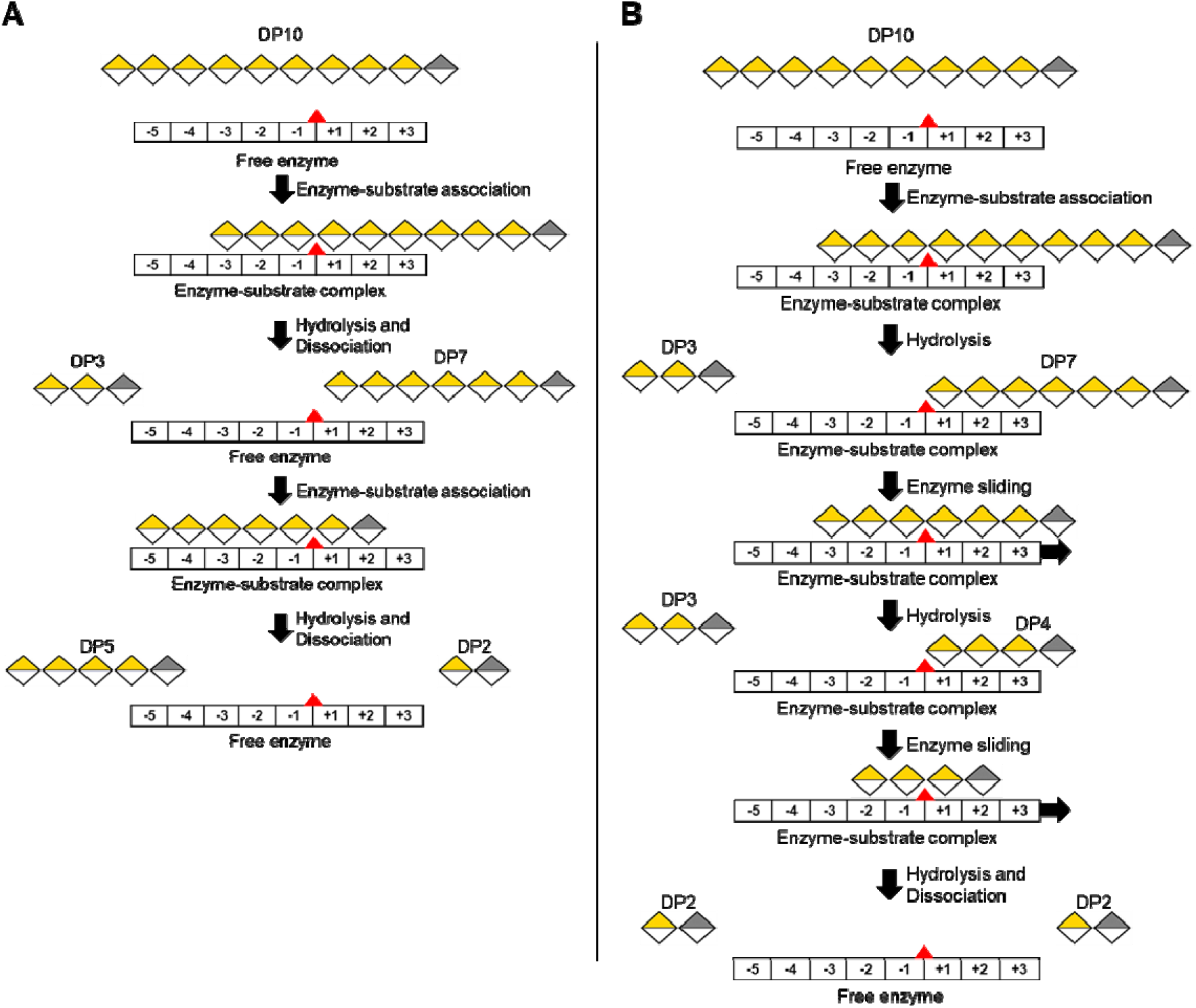
Model of PGLR and ADPG2 processivity. (A) PGLR shows low processive dynamics where enzyme-substrate association is followed by hydrolysis and dissociation of the substrate from the enzyme. This low processivity produce OGs of variable DPs. B) ADPG2 sliding motion after forming enzyme-substrate complex allows multiple substrate hydrolysis while staying attached to the substrate showing highly processive dynamics. Processive enzymes can produce small DP OGs. Galacturonic acid are yellow colored. Galacturonic acid reducing end is grey colored. PG subsites are indicated by numbers. Red triangle represents the hydrolysis site.

Our work demonstrates that albeit having a highly conserved structural fold, subtle differences in PG structures translate into differences in enzymes’ dynamics, substrate specificities, kinetics, leading to distinct processivities that translate into distinct impacts on plant development. This shows the extent by which, among the multigenic family, each of the isoforms has peculiar specificities that would be required, at the cell wall, to control temporally and spatially pectin structure. This further highlight that, for this class of enzymes, the gene redundancy at the genome level is unlikely to reflect redundant biochemical specificities. Our study now paves the way for a better understanding of how PG’s processivities can control polysaccharides chemistry and mechanical properties *in muro*.

## Material and methods

### Sequences retrieval and analysis

*Arabidopsis thaliana* PGLR (At5g14650) and ADPG2 (At2g48150) coding sequences were retrieved from publicly available genome database TAIR (https://www.arabidopsis.org/). The presence of putative signal peptide was predicted using SignalP-5.0 Server (http://www.cbs.dtu.dk/services/SignalP/). Glycosylation sites were predicted using NetNGlyc 1.0 Server (http://www.cbs.dtu.dk/services/NetNGlyc/). Sequence alignments were performed using MEGA and Clustal Omega multiple sequence alignment programs (41).

### Cloning, heterologous expression and purification of PGLR and ADPG2

PGLR was previously expressed in the yeast *Pichia pastoris* and biochemically characterized (8). PGLR mutants were created using cDNA and specific primers carrying mutations (SI Appendix, Table S1). At2g41850 (ADPG2) coding sequence was codon-optimized for *Pichia pastoris* expression. Cloning and protein expression was done as previously described (8, 42).

### PGLR and ADPG2 enzyme analysis

Bradford method was used to determine the protein concentration, with bovine serum albumin (A7906, Sigma) as a standard. Deglycosylation was performed using Peptide-N-Glycosidase F (PNGase F) at 37 °C for one hour according to the supplier’s protocol (New England Biolabs, Hitchin, UK). Enzyme purity and molecular weight were estimated by 12 % SDS-PAGE using mini-PROTEAN 3 system (BioRad, Hercules, California, United States). Gels were stained using PageBlue Protein Staining Solution (Thermo Fisher Scientific) according to the manufacturer’s protocol.

### PGLR and ADPG2 biochemical characterization

The substrate specificity of PGLR and ADPG2 were determined with the DNS method as previously described (8, 42). PolyGalacturonic Acid (PGA, 81325, Sigma); Citrus pectin with DM 20-34% (P9311, Sigma), DM 55-70% (P9436, Sigma) were used as substrates. Results were expressed as nmol of GalA.min^−1^.μg^−1^ of proteins. The optimum temperature was determined by incubating the enzymatic reaction between 25 and 60°C during 60 min using PGA (0.4%, w/v) at pH5. The pH optimum was determined between pH 4 and 7 using 100 mM sodium acetate buffer (pH 3 to 5) and phosphate citrate buffer (pH 6 to 8) and 0.4 % (w/v) PGA as a substrate. The PGLR and ADPG2 kinetic parameters were calculated using GraphPad Prism8 (version 8.4.2.) with PGA as a substrate. The reactions were performed using 1 to 8 mg.ml^−1^ PGA concentrations during 10 min at 25°C in 50 mM sodium acetate (pH5). All experiments were realized in triplicate.

### Digestion of cell wall pectins and released OGs profiling

OGs released after digestions by recombinant PGLR and ADPG2 were identified as described (31). Briefly, PGA (81325, Sigma) or citrus pectin with DM 24-30% (P9311, Sigma) or OGs DP12DM5 (degree of polymerization centered on 12 and average DM of 5%) were prepared at 0.4 % (w/v) final concentration diluted in 100 mM ammonium acetate buffer (pH 5) and incubated with either PGLR and ADPG2 at 0.03 μg.μL^−1^. Non-digested pectins were pelleted by centrifugation and the supernatant dried in speed vacuum concentrator (Concentrator plus, Eppendorf, Hamburg, Germany). The same procedure was applied for pectins from roots of Arabidopsis seedlings that were grown for 7 days at 21 °C, 16 h/8 h light/dark photoperiod. Roots were cut, incubated in ethanol 100 % (w/v) for 24 h, washed two times 5 min with acetone 100 % (w/v) and left to dry 24 h. Thirty roots per replicate were rehydrated in 150 μL 100 mM ammonium acetate pH 5 during 2 h on room temperature and digested with PGLR and ADPG2 at 0.02 μg.μL^−1^ on average, using the above-mentioned protocol. Separation of OGs was done using an ACQUITY UPLC Protein BEH SEC column (125Å, 1.7 μm, 4.6 mm x 300 mm), and the analysis was done as described (42).

### Microscale thermophoresis

Molecular interactions between PGLRs (WT and mutants) and DP12DM5 was done using Microscale thermophoresis (MST) approach as described with some modifications (43). Briefly, PGLRs were labelled with monolith protein labelling kit blue NHS amine reactive (Lys, NanoTemper, catalog no. MO-L003) and conserved in MST buffer (50 mM Tris pH 7.4, 150 mM NaCl, 10 mM MgCl2, 0.05 % tween-20). For all experiments, constant final concentration of labelled PGLRs was 1650 nM. Mix of OGs centred on DP12DM5 was prepared at 14028 nM concentration in MST buffer/dH_2_O in 1:1 ratio. For all experiments, a constant concentration of labelled PGLRs was titrated with decreasing concentrations of non-labelled DP12DM5 from 7014 to 0.214 nM. The resulting mixtures were loaded into a Monolith NT.115 series standard capillaries (NanoTemper, catalogue no. MO-K002). Thermophoresis experiments were performed with 40% of MST power and 20% of LED power for fluorescence acquisition.

### Time-resolved molecular dynamics measurements

PGLR and ADPG2 (used as ligands) were immobilized on an electro-switchable DNA biochip MPC-48-2-R1-S placed into a biosensor analyzer switchSENSE® DRX (Dynamic Biosensors GmbH, Planegg, Germany). For that, a covalent conjugate between PGLR or ADPG2 and a 48mer ssDNA was first prepared with the amine coupling kit supplied by Dynamic Biosensors and purified by anion-exchange chromatography onto a proFIRE® system (Dynamic Biosensors), then hybridized with a complementary ssDNA attached on the surface of the biochip and carrying a Cy5 fluorescent probe at its free extremity. When analytes injected in the microfluidic system bind to the oscillating dsDNA nanolevers, the nanolever movement is altered by the additional friction imposed. Kinetic measurements for 2 min (association) and for 5 min (dissociation) were performed in 5 mM sodium acetate buffer, pH 5.5, with a flow rate of 100 μl.min^−1^ at 25°C with different concentrations of various analytes: PGA (81325, Sigma), citrus pectin with DM 24-30% (P9311, Sigma) and pool of OGs centred on DP12DM5, DP12DM30 and DP12DM60 at 25, 50 and 100 μM. The fluorescence traces were analysed with the switchANALYSIS® software (V1.9.0.33, Dynamic Biosensors). The association and dissociation rates (k_on_ and k_off_), dissociation constant (K_D_ = k_off_/k_on_) and the error values were derived from a global single exponential fit model, upon double referencing correction (blank and real-time) (44). The experiments were performed in three replicates.

### Intrinsic processivity calculations

The intrinsic processivity potential (P^Intr^), a parameter corresponding to the number of consecutive catalytic steps before dissociation from the substrate was used as a measure of the processivities of PGLR and ADPG2 as described in Horn et al. (32). The calculation of P^Intr^ is given in the Eq. 1.

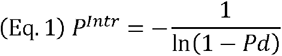

The dissociation probability (P_d_) is expressed as a rate constant for two processes; (i) the turnover number (k_cat_) and (ii) the enzyme–substrate complex dissociation constant (k_off_). P_d_ is related to k_cat_ and k_off_ according to Eq. 2. In the case of processive enzymes P_d_ « 1.

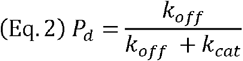

The turnover number (k_cat_) was calculated using GraphPad Prism8 (version 8.4.2.) by fitting the non-linear regression curve following the Eq. 3, where Y is enzyme velocity, X is the substrate concentration, Km is the Michaelis-Menten constant in the same units as X and Et is the concentration of enzyme catalytic sites, 0.02307 and 0.001944 nM for PGLR and ADPG2, respectively.

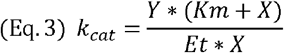

### Crystallization of proteins

PGLR and ADPG2 were concentrated at 10 mg.ml^−1^. Crystallization was performed using the sitting-drop vapour-diffusion method. Crystallisation conditions were screened using a mosquito robot (TTP Labtech) and the PACT premier plate (Molecular dimensions, Sheffield, UK). PGLR and ADPG2 (100 nL) were mixed with an equal volume of precipitant (1:1). The crystals that resulted in best diffraction data were obtained with 0.2 M sodium fluoride, 0.1 M bis-tris propane pH 8.5, 20 % w/v PEG 3350 (H1 condition, PACT premier plate) for PGLR and 0.2 M sodium malonate dibasic monohydrate, 20 % w/v PEG 3350 (E12 condition, PACT premier plate) for ADPG2. Crystals for PGLR and ADPG2 formed after 6 and 2 months, respectively. Scale-up of the best condition was realized by mixing 1 μl of the best precipitant condition with 1 μl of the enzyme in the hanging drop vapor-diffusion method.

### X-ray data collection and processing

Crystals were mixed with precipitation solution and PEG 3350 (35% w/v) before mounting in a loop and flash cooling in liquid nitrogen. The diffraction data were collected at PROXIMA-1 beamline (Synchrotron Soleil, Saint Aubin, France), at a temperature of -173°C using a PILATUS 6M end EIGER 16M detector (Dectris). Data were collected using X-rays with wavelength of 0. 978564 Å. For PGLR, three data sets were collected from the same crystal at 1.3 Å resolution. Intensities were integrated, scaled and merged using XDS (45) and XSCALE (46). For ADPG2, one data set was collected at 2.03 Å resolution. Intensities were processed using XDS (45). PGLR crystal belonged to triclinic space group P1 with one molecule in asymmetric unit, while ADPG2 belongs to orthorhombic space group P2_1_2_1_2_1_ with two molecules in asymmetric units.

### Structure solution and refinement and analysis

For PGLR and ADPG2 structure and function prediction I-TASSER prediction software was used (47). ColabFold was used for AtPGIP1 and AtPGIP2 models (48). The structure of PGLR was solved by molecular replacement using *Phaser*(49). The data were phased using rhamnogalacturonase A (PDB: 1RMG, Uniprot: Q00001) as a search model (23). Model was built using *Autobuild* and refined using *Refine* from PHENIX suite (50). The model was iteratively improved with *Coot* (51) and *Refine*. ADPG2 structure was solved by molecular replacement using PGLR as a starting model and the above-mentioned iterative procedure. The final structure for PGLR and ADPG2 have been deposited in the Protein Data Bank as entries 7B7A and 7B8B, respectively. UCSF Chimera was used for creation of graphics (52).

### Modelling and molecular dynamics simulations

Molecular dynamics (MD) simulations were carried out on both the WT PGLR and ADPG2 proteins in complex with fully de-methylesterified decasaccharides, as well as partially methylesterified decasaccharides. Additionally, PGLR mutants H196K and H237K, modelled from the resolved X-ray crystal structures using PyMOL, were also simulated, in complex with fully de-methylesterified decasaccharides (53).

Molecular topologies of the complexes were created according to the parameters of the AMBER14SB_parmbsc1 forcefield (54). The complexes were placed in cubic boxes with a solute-box distance of 1.0 nm and solvated with water molecules parameterised according to the TIP3P water model (55). To neutralise the system’s net charge and reach a salt concentration of 0.165 M, Na^+^ and Cl^−^ ions were added before energy-minimisation was performed.

The systems were then energy minimized, to resolve clashes between particles using a steep-descent algorithm with a step size of 0.01, considering convergence when the particle-particle force was 1000 kJ mol^−1^ nm^−1^. Particle-particle forces were computing considering van der Waals and electrostatic interactions occurring up to 1.0 nm, treating long-range electrostatics in the Fourier space using the Particle Mesh Ewald (PME) summation method.

After minimization, solvent equilibration was achieved in two stages to reach constant temperature and pressure. The first stage was performed in the nVT ensemble while the second in the nPT ensemble. Solvent equilibration through the nVT ensemble was carried out for 1 ns, with the equation of motion integrated with a time step of 2 fs, targeting a reference temperature of 310.15 K coupled every 0.1 ps using the V-rescale thermostat (56).

In this step, each particle in the system was assigned random velocities based on the Maxwell-Boltzmann distribution (57) obtained at 310.15 K. Equilibration of the solvent through the nPT ensemble was then carried out for 1 ns starting from the last step (coordinates and velocities) of the previous equilibration, at a reference temperature of 310.15 K, coupled every 0.1 ps using the V-rescale thermostat (56). In this step, pressure coupling was conducted at 1 bar, with pressure coupled isotropically every 2.0 ps using the Parrinello-Rahman barostat(58). Particle-particle interactions were calculated by building pair lists using the Verlet scheme. A cutoff of 1.0 nm was used to compute short-range van der Waals and electrostatic interactions sampled via a Coulomb potential. The Particle Mesh Ewald (PME) algorithm (59), with a Fourier grid spacing of 0.16 and a cubic B-spline interpolation level of 4, was used to compute, in the Fourier space, long-range electrostatic interactions past the cutoff.

Simulations were then performed on both in-house machines and on NeSI’s (New Zealand eScience Infrastructure) high performance cluster, Mahuika, using GROMACS (Groningen MAchine for Chemical Simulation) version 2020.5 (60). For each of the 6 complexes, simulations were run for 200 ns using a time step of 2 fs and replicated 5 times for a total simulation time of 1 μs per complex. Each replicate differed in terms of the random sets of particle velocities generated through the nVT ensemble. Molecular dynamics trajectories were recorded every 10 ps. For analysis, the first 50 ns of each production run were considered equilibration time and discarded.

Analyses were conducted using in-house Python 3 scripts implemented Jupyter notebooks (61). Porcupine plots were created using data from a normalised principal component analysis calculated using GROMACS. Figures were created and rendered with Matplotlib (62), VMD (Visual Molecular Dynamics) (63) and UCSF Chimera (52).

### Poisson-Boltzmann calculations of electrostatic potentials

The protonation states of each amino acid were assigned according to the pKa curves calculated at pH = 4 for PGLR and pH = 5 for ADPG2, using the PROPKA software (64). Atomic charges and radii for the protein atoms were assigned using the PDB2PQR software (65) according to the parameters of the AMBER14SB_parmbsc1 forcefield (54), while atomic charges and radii for the sugar atoms were obtained from our previous work (66). The surface electrostatic potentials for WT PGLR and ADPG2 were then calculated solving the non-linearized form of the Poisson-Boltzmann equation through the APBS (Adaptive Poisson–Boltzmann Solver) software on a cubic grid composed of 193 grid points across the x-, y-and z-directions (67). These calculations followed a stepwise approach where the Poisson-Boltzmann equation is first solved on a coarse mesh grid with a length of 155 Å and a spacing of 0.8 Å; then on a fine mesh grid with a length of 125 Å and a spacing of 0.64 Å. Calculations were solved considering a temperature of 218.15 K with a mobile ionic charge of +/-1 e_c_, an ionic concentration of 0.165 M and an ionic radius of 2.0 Å. The protein dielectric constant was set at 4.0 and the solvent dielectric constant was set to 78.54. The protein surface electrostatic potentials were then visualised and coloured on the protein’s molecular surface using VMD (63).

### Exogenous application of enzymes on Arabidopsis seedlings

20-30 sterile seeds of *Arabidopsis thaliana* EGFP-LTI6b(34) plasma membrane marker-lines were sowed in 400 μl liquid Arabidopsis Murashige and Skoog medium (Sucrose, MES (Duchefa monohydrate), in 24 well-plates (68). After 48 hours stratification, plates were placed in growth chamber under long day conditions (16 hours photoperiod, 120 μmol.m^−2^.s^−1^, 21°C). After 6 days, cultures were supplemented with 0.051 μg/μl and 0.015 μg/μl filter-sterilized PGLR and ADPG2, respectively using 0.2 μm PES filter (Whatman TM Puradisc TM 13 mm) in a volume of liquid MS medium of 200 μl to reach iso-activity. Plantlets were allowed to grown for another 1 day (T1) or 3 days (T3). Negative controls correspond to 6-, 7-or 9-days cultures with buffer only (T0 ØEnz, T1 ØEnz and T3 ØEnz, respectively). For each of these conditions, measurements of primary root lengths were done using ImageJ software with NeuronJ plugin. For each condition, 30-40 plants were measured. For cell lengths determination, approximately 1 mm from the tip of the root of 3 to 7 plants were photographed under a stereomicroscope (ZEISS SteREO Discovery.V20). Images were assembled using MosaicJ plugin from Image J. The length of the firsts 50 rhizodermal cells, starting from the first cell of the columella, were measured using image J software with NeuronJ plugin. Phenotypical observations where performed following ruthenium red staining (0.05 % (w/v) in water, Sigma-Aldrich R-2751) under binocular microscope (Leica EZ4).

## Acknowledgements

This work was supported by a grant from the Agence Nationale de la Recherche (ANR-17-CE20-0023) and by the Conseil Regional Hauts-de-France and the FEDER (Fonds Européen de Développement Régional) through a PhD grant awarded to J.S. We wish to thank Pierre Legrand and all the staff at Proxima1 beamline (Synchrotron SOLEIL, Gif sur Yvette, France) for X-ray diffraction and data collection. The technical assistance of Maša Boras, a former master student is gratefully acknowledged.

