## Supplemental figures for "Differences in the structure of plant polygalacturonases specify enzymes’ dynamics and processivities to fine-tune cell wall pectins"

^1^UMRT INRAE 1158 BioEcoAgro – BIOPI Biologie des Plantes et Innovation, Université de Picardie, 33 Rue St Leu, 80039 Amiens, France. ^2^School of Chemical Sciences, The University of Auckland, Private Bag 92019, Auckland 1142, New Zealand. ^3^UMR 8576 Unité de Glycobiologie Structurale et Fonctionnelle (UGSF), 50 Avenue de Halley, 59658 Villeneuve d’Ascq, France. ^4^Université Paris-Saclay, INRAE, AgroParisTech, Institut Jean-Pierre Bourgin (IJPB), 78000, Versailles, France ^5^Plateforme Analytique, Université de Picardie, 33, Rue St Leu, 80039 Amiens, France. ^6^INRAE, UR 1268 Biopolymers, Interactions Assemblies, CS 71627, 44316 Nantes Cedex 3, France. ^7^Université de Lorraine, INRAE, IAM, F-54000 Nancy, France.

#: Contributed equally as first authors, §: Contributed equally as last authors

**Corresponding authors:**

Jérôme Pelloux

Davide Mercadante

Fabien Sénéchal

**This PDF file includes:**

Figures S1 to S12

Tables S1

Legends for Datasets S1

SI References

**Other supporting materials for this manuscript include the following:**

Datasets S1

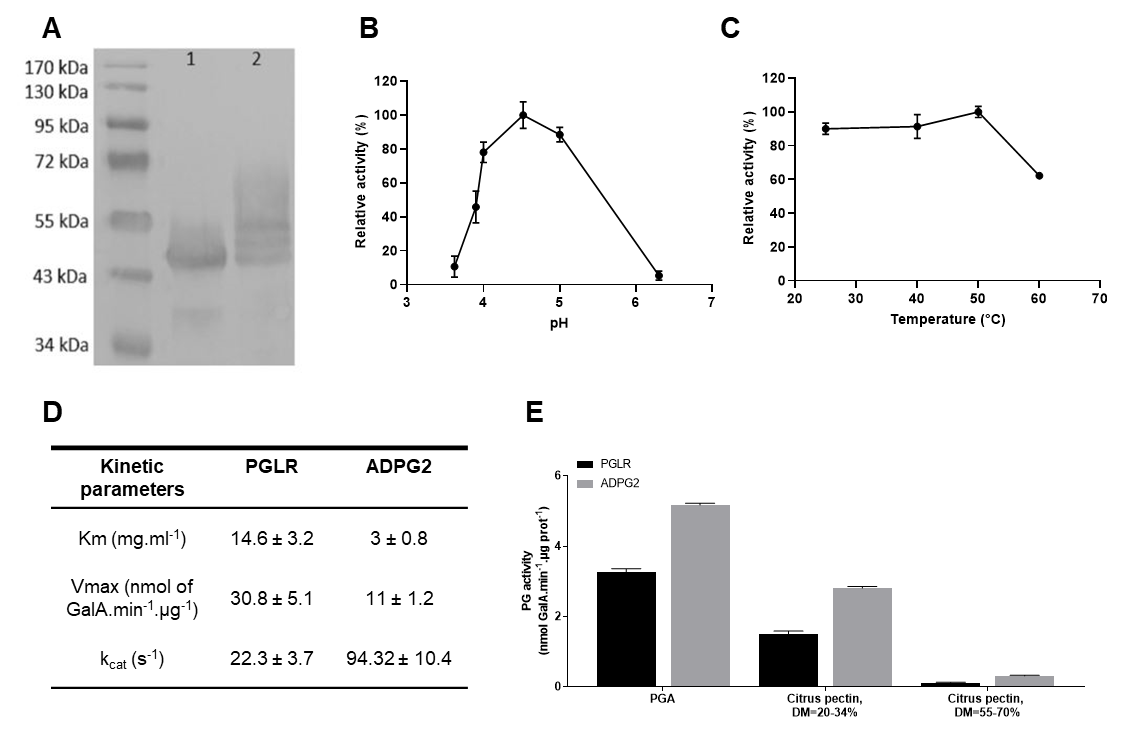

**Fig. S1: Purification and biochemical characterization of ADPG2**

A) Western blot analysis of ADPG2 with anti-His antibodies on de-glycosylated form of ADPG2 obtained after digestion by PNGase F (1) and non-digested native sample (2). B) pH-dependence of ADPG2 activity. The activities were measured after 1 hour of incubation with PGA at 25°C at various pHs. C) Temperature-dependence of ADPG2 activity. The activities were measured after 1 hour of incubation with PGA at pH 5.2 at various temperature. D) Determination of Km, Vmax and k_cat_ for ADPG2 and PGLR. Activity was assessed using various concentrations of polygalacturonic acid (PGA) at 25°C and pH 5.2 using the 3,5-dinitrosalicylic acid method. E) Substrate specificity of PGLR and ADPG2. Activity was measured at 50°C and pH 5.2 during 1 hour using substrates of increasing degrees of methylesterification.

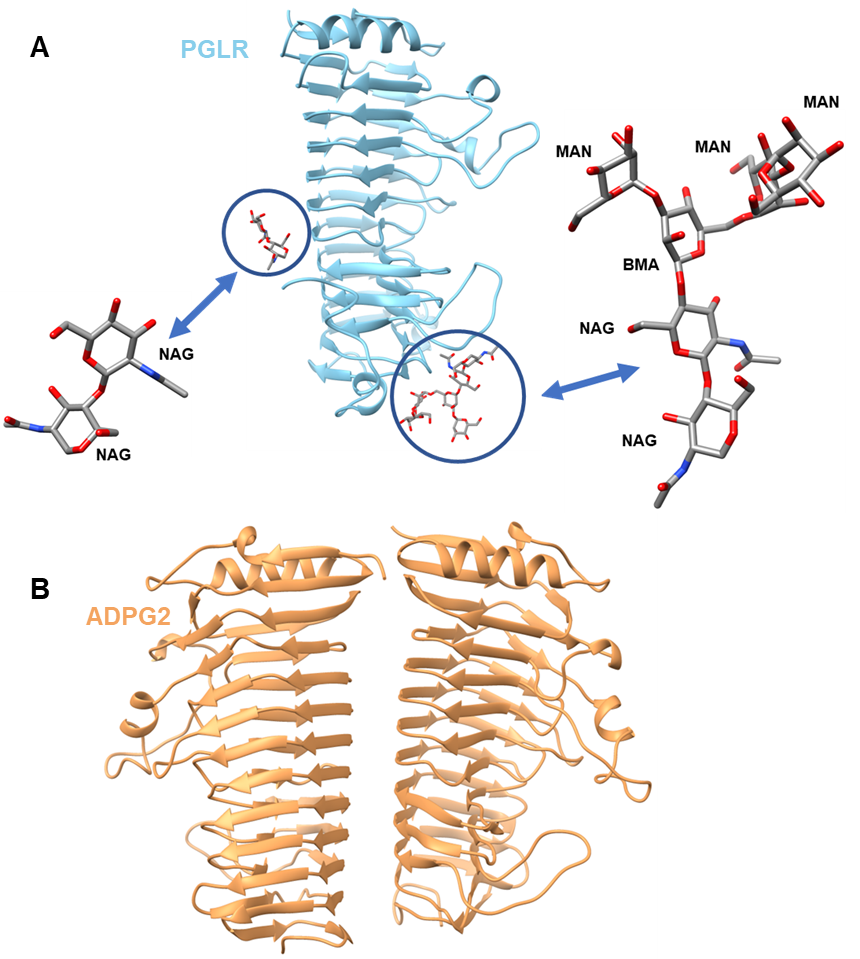

**Fig. S2. Crystallised PGLR and ADPG2 in asymmetric unit and glycosylation sites**

A) Ribbon diagram of the PGLR structure containing 1 molecule in the asymmetric unit. PGLR harboured two N-glycosylation sites: N255- linked NAG-NAG and N313-linked NAG–NAG–BMA–MAN–MAN–MAN. NAG; N-acetylglucosamine, BMA; β-mannose, MAN α-mannose. B) Ribbon diagram of the ADPG2 structure containing 2 molecules in the asymmetric unit.

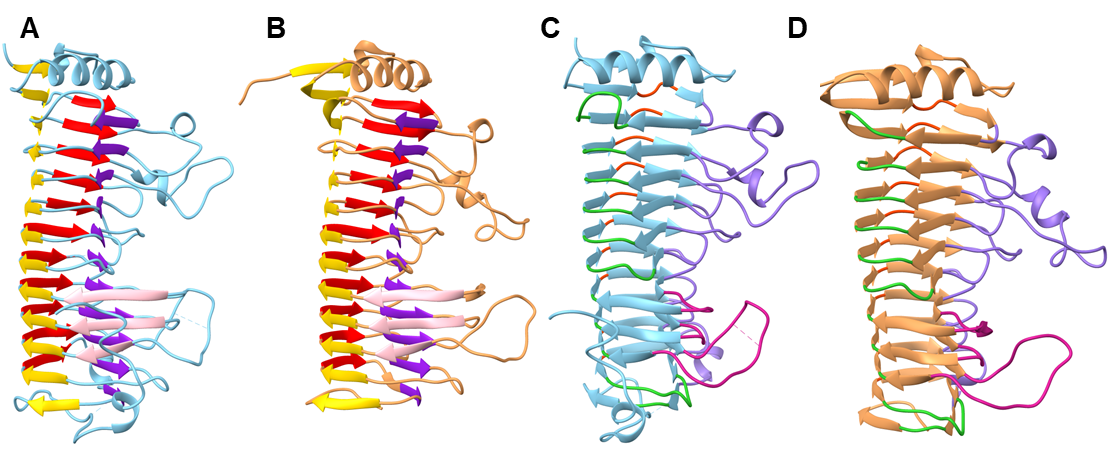

**Fig. S3. PGLR and ADPG2 represent right-handed parallel β-helical structure**

Ribbon structure representing β-sheets (PB1-purple, PB1a-pink, PB2-yellow and PB3-red) for PGLR (A) and ADPG2 (B). Ribbon structure representing T-turns (T1-lime green, T1a-violet red, T2- orange red, T3 medium purple) for PGLR (C) and for ADPG2 (D). β-strands and T-turns are named according to Petersen et al. 1997 (1).

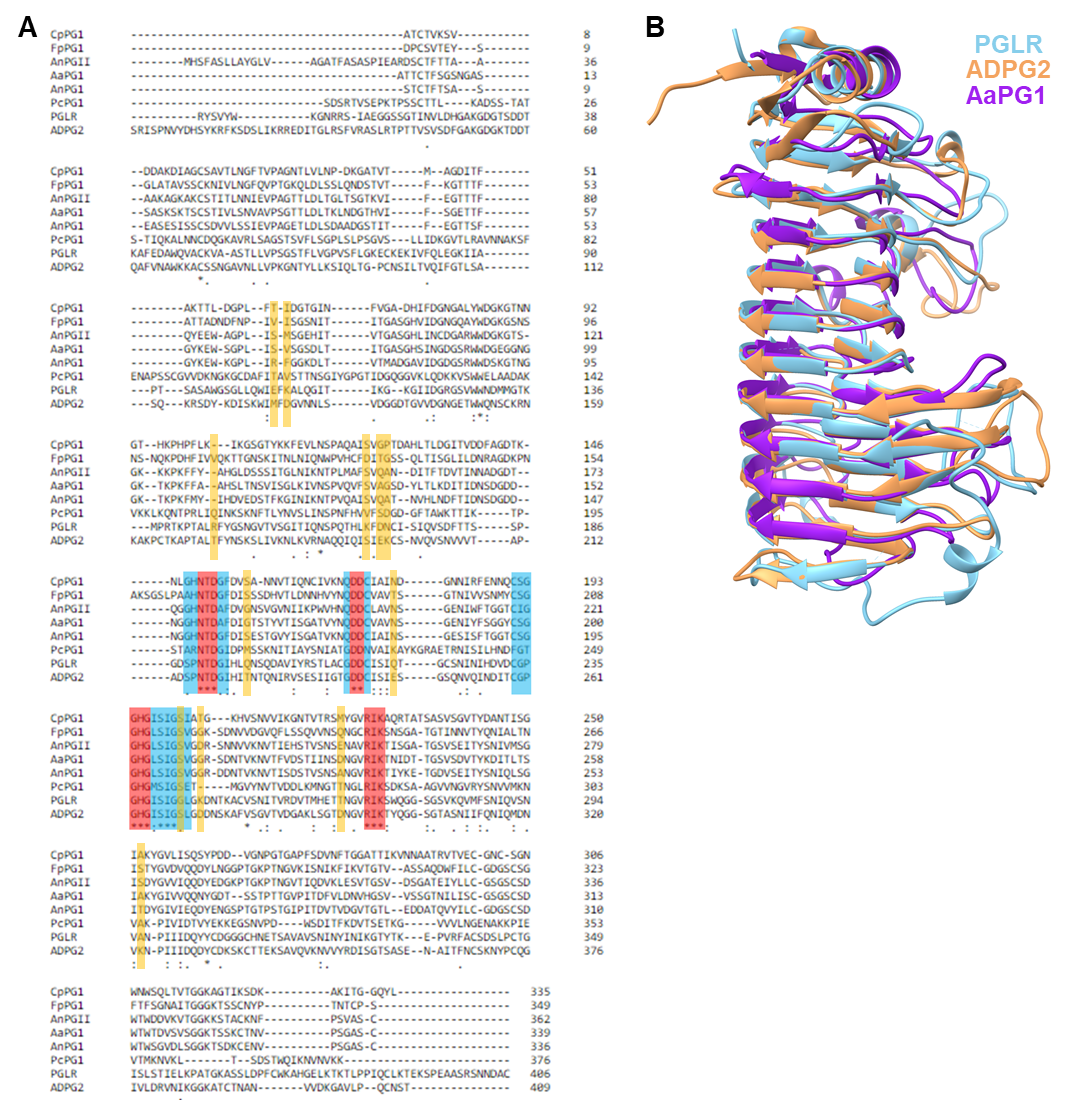

**Fig. S4. PGLR and ADPG2 sequence and structure identity with selected fungal enzymes**

A) Sequence alignment of PGLR and ADPG2 with characterized fungal PGs. Selected PGs; *Pectobacterium* *carotovorum* PG1 (PcPG1, PDB: 1BHE), *Aspergillus niger* PGI (AnPGI, PDB: 1NHC) and PGII (AnPGII PDB: 1CZF), *Fusarium phyllophilum* PG1 (FpPG1, 1HG8), *Aspergillus aculeatus* (AaPG1, PDB: 1IB4) and *Chondrostereum purpureum* (CpPG1, PDB: 1KCD). The aa of the active are red-boxed, the conserved aa are blue-boxed and binding site aa that differ between the PGLA and ADPG2 are yellow boxed (2). The alignment was performed using ClustalO. B) Superimposition of PGLR, ADPG2 and AaPG1 structures (3).

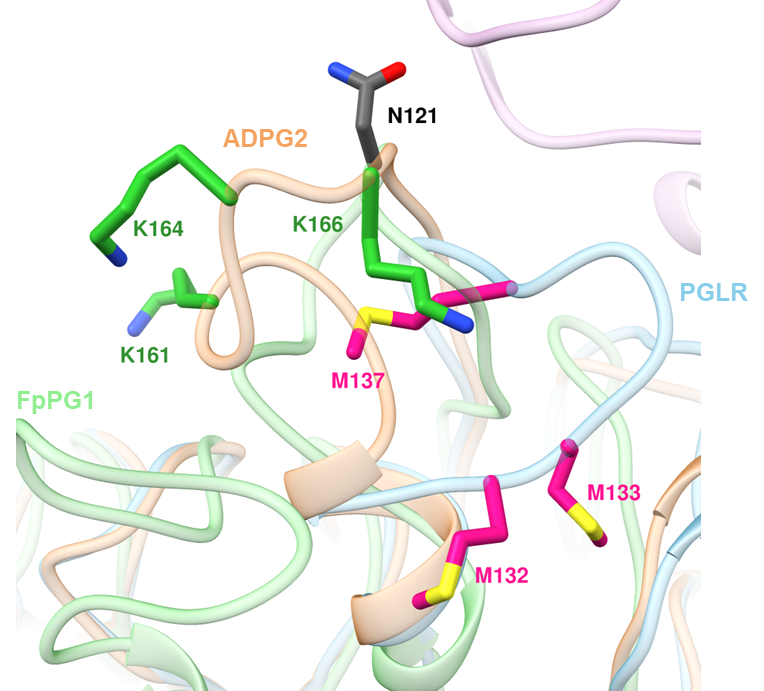

**Fig. S5. PGLR and ADPG2 N-terminal loops**

In PvPGIP2-FpPG1 interaction N-terminal loop with N121 play a key role in PG-PGIP interaction (FpPG1 aa in grey). In PGLR (blue) this loop is rich in methionine (pink) while ADPG2 loop (in brown) is rich in lysine (green) residues. PvPGIP2 is plum-colored.

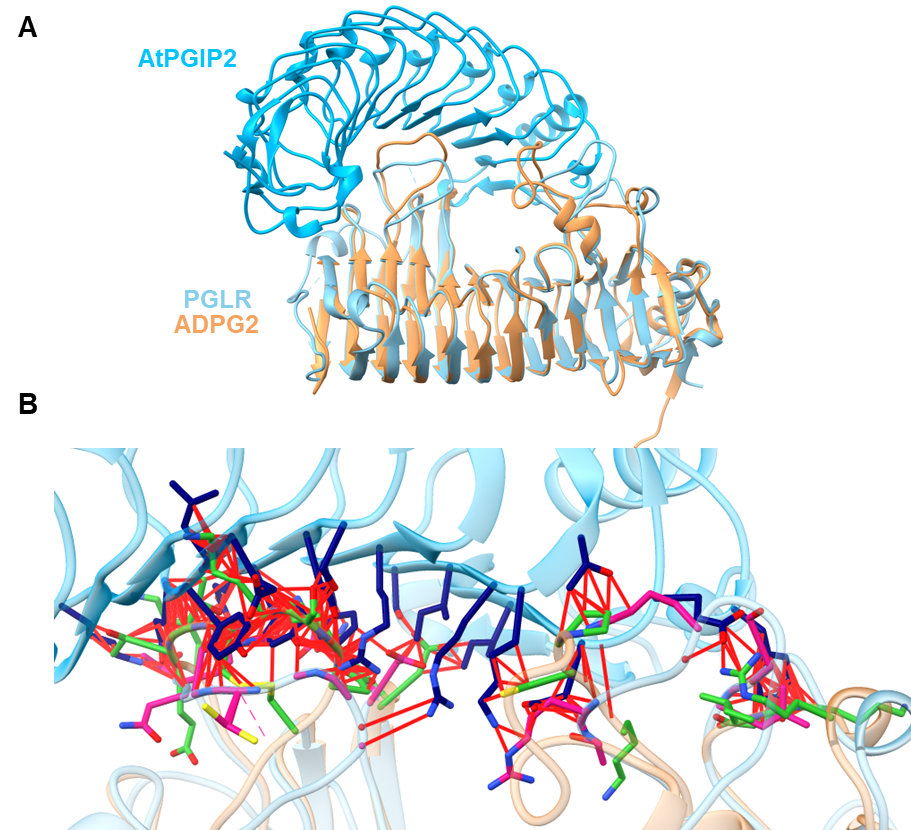

**Fig. S6. Structural determinants of the absence of interaction between AtPGIP2 and PGLR-ADPG2**

A) Ribbon representation of AtPGIP2 (dark blue) in interaction with PGLR (blue) and ADPG2 (brown). B) Interaction of AtPGIP2 with PGLR and ADPG2. The model of AtPGIP2 was superimposed onto PvPGIP2. Amino acids of AtPGIP2 (navy blue), PGLR (pink) and ADPG2 (green) included in clashes closer than 0.6 Å are shown. The red lines represent atoms overlap of minimum 0.6 Å.

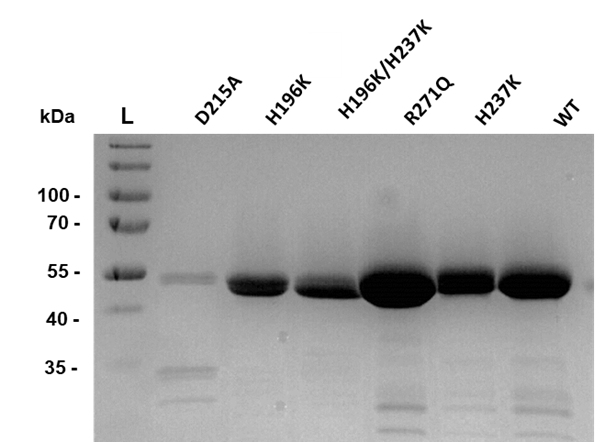

**Fig. S7. SDS-PAGE representing the wild type and mutants of PGLR**

PGLR and its mutants were purified with His-tag using 1 mL Ni-NTA colon. Proteins were resolved on a 12% polyacrylamide gel and were stained by Coomassie blue. L-ladder.

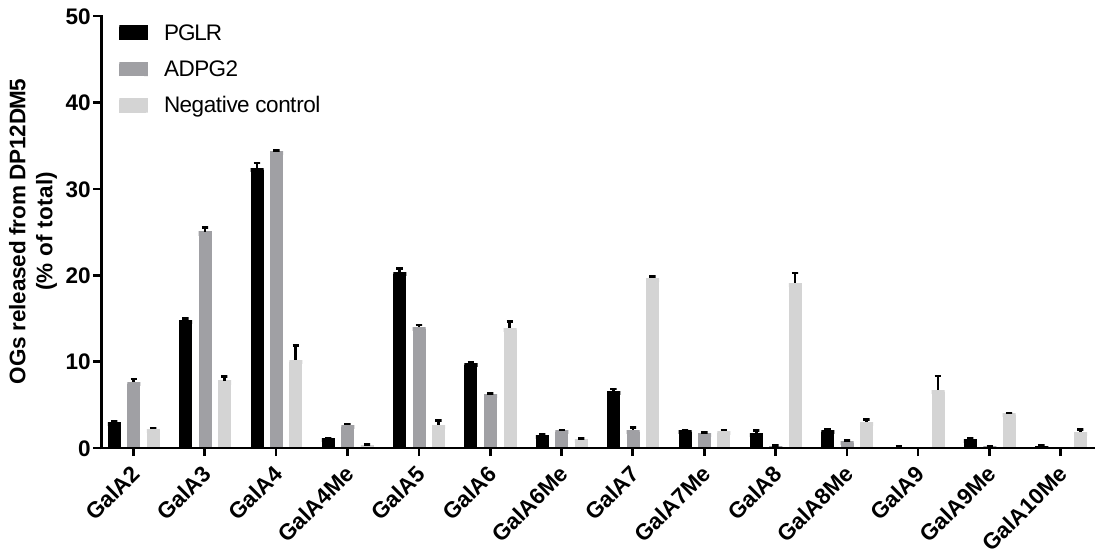

**Fig. S8. Oligogalacturonides produced by PGLR and ADPG2 from pectins of DP12DM5**

Oligoprofiling of OGs after over-night digestion of DP12DM5 (degree of polymerization centered on 12 and degree of methylesterification centred on 5) pectins by PGLR and ADPG2 at 40°C, pH5.Negative control: undigested DP12DM5 pectins.

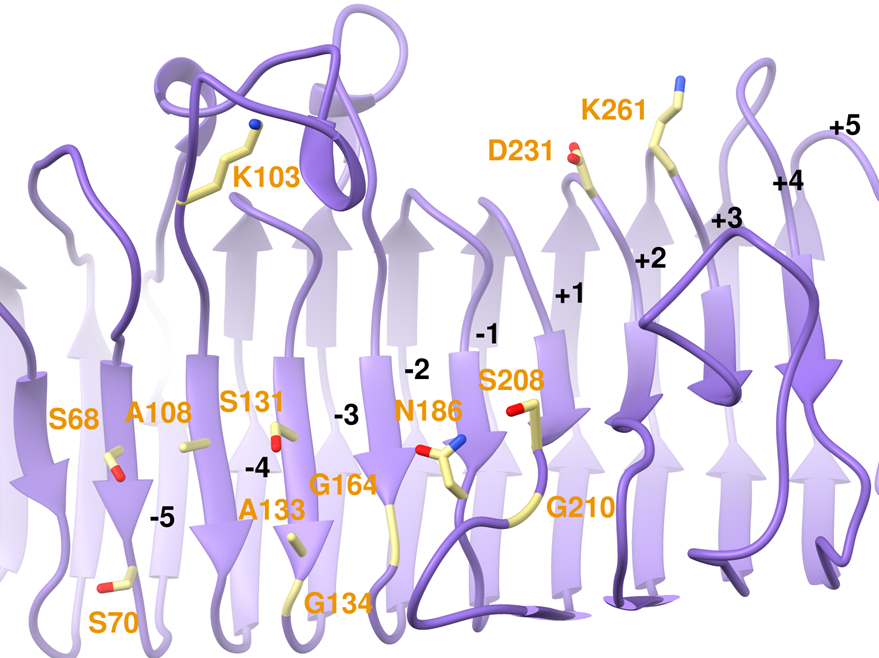

**Fig. S9. Structure of subsites of AaPG1**

Structure of the -5/+5 subsites of AaPG1 (purple with khaki labelled aa).

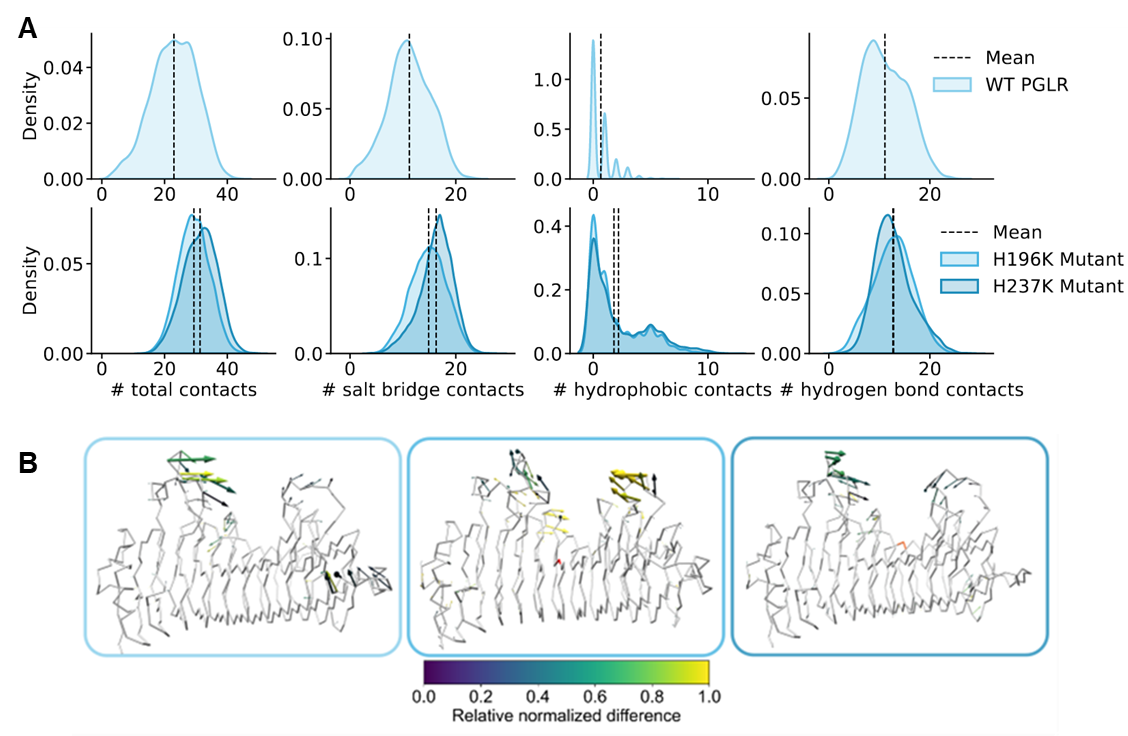

**Fig. S10. PGLR H196K and H237K mutants contact calculations**

A) Probability density distributions for the number of contacts that occur between WT PGLR (light blue), H196K (medium blue) and H237K mutants (dark blue) and their fully de-methylesterified decasaccharides, within a cutoff of 4.0 Å. Black dashed lines indicate mean values. B) Normalized porcupine plots illustrating the largest motions of protein alpha-carbon atoms represented by the first eigenvector of a principal component analysis, for WT PGLR (light blue; left), H196K (medium blue; middle) and H237K (dark blue; right); complexed to fully de-methylesterified decasaccharides (pattern 1). Arrows indicate direction and relative normalized magnitude of movement, from 0 (dark blue) to 1 (yellow).

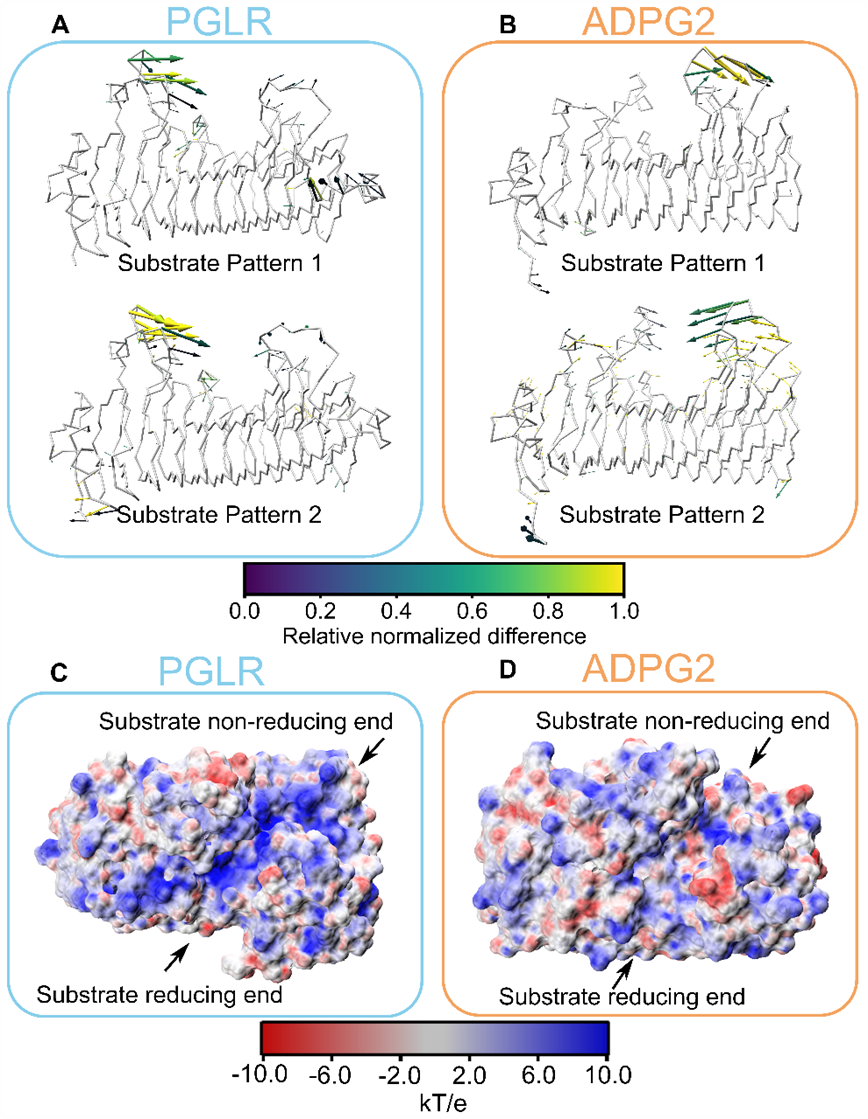

**Fig. S11.** **Porcupine plots and surface electrostatic potential of PGLR and ADPG2**

A) Normalized porcupine plots illustrating the largest motions of protein alpha-carbon atoms represented by the first eigenvector of a principal component analysis, for WT PGLR with a fully demethylesterified decsaccharide (pattern 1) (top) and a 60% methylesterified decsaccharide (pattern 2, bottom). Arrows indicate direction and relative normalized magnitude of movement, from 0 (dark blue) to 1 (yellow). B) Normalized porcupine plots illustrating the largest motions of protein alpha-carbon atoms represented by the first eigenvector of a principal component analysis, for WT ADPG2 with a fully demethylesterified decsaccharide (pattern 1, top) and a 60% methylesterified decsaccharide (pattern 2, bottom). Arrows indicate direction and relative normalized magnitude of movement, from 0 (dark blue) to 1 (yellow). C) WT PGLR protein surface electrostatic potential projected on the protein’s molecular surface, coloured from -10 kT/e (red) to 10 kT/e (blue). The non-reducing (subsite -5) and reducing (subsite +5) ends are labelled as appropriate. D) WT ADPG2 protein surface electrostatic potential projected on the protein’s molecular surface, coloured from -10 kT/e (red) to 10 kT/e (blue). The non-reducing (subsite -5) and reducing (subsite +5) ends are labelled as appropriate.

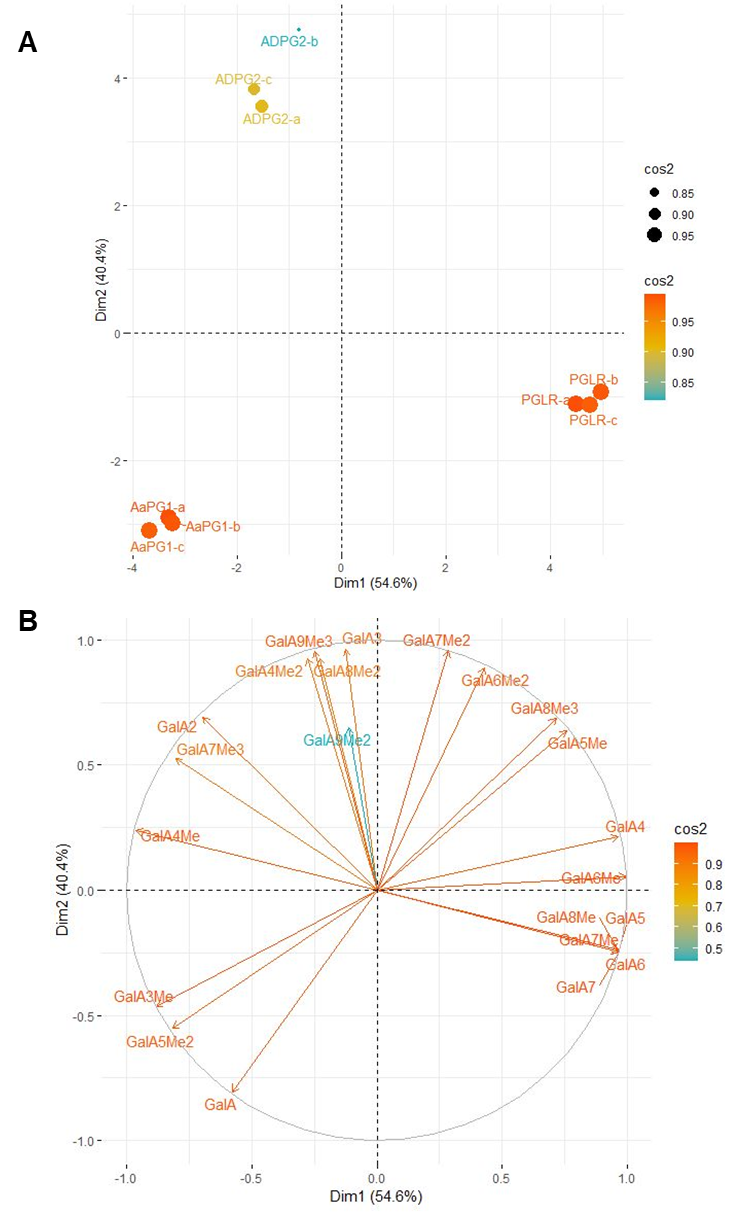

**Fig S12. PCA of OGs produced by PGLR, ADPG2 and AaPG1**

A) Score plot of Principal Component Analysis (PCA) of oligogalaturonides released from pectins DM 20-34% by PGLR, ADPG2 and AaPG1. a, b, c represents biological repetitions. B) Loading plot. The oligogalacturonides released after overnight digestions of pectins DM 20-34% by PGLR, ADPG2 and AaPG1 were analysed by PCA using R-package (FactoMineR and Factoextra).

**Table S1. Primers for cloning mutated forms of PGLR and ADPG2 into pPICzαB expression vectors.**

Table with primers used for the cloning of coding sequences into pPICzαB. Restriction enzymes sites for EcoRI, NotI are underlined, added bases are written in *italics*. Mutation bases are **bolded**.

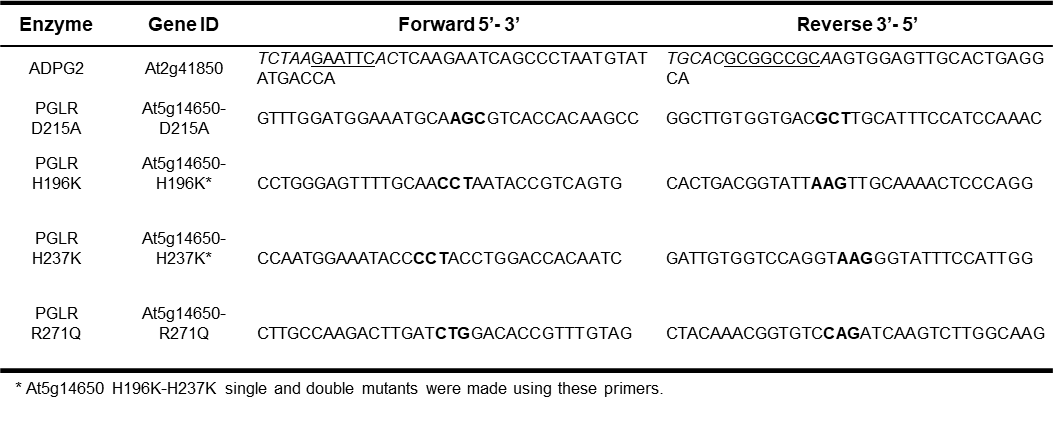

**Dataset S1 (separate file).** AtPGIP1 and AtPGIP2 contact analysis with PGLR and ADPG2

Contact analysis was done using Chimera. All contact between the atoms closer then 0.6 Å are listed.
